# Anti-HIV-1 B cell antigen receptor signaling and structure

**DOI:** 10.1101/2024.11.15.623645

**Authors:** Bhishem Thakur, Jahangir Alam, Kenneth Cronin, Parth Patel, Kara Anasti, Advaiti Pai Kane, Alamgir Hossain, Hung Do, Katayoun Mansouri, Taylor N. Spence, Robert J. Edwards, Katarzyna Janowska, Muralikrishna Lella, Aaron Cook, Kevin Saunders, Sandrasegaram Gnanakaran, Barton F. Haynes, Priyamvada Acharya, S. Munir Alam

## Abstract

The B cell antigen receptor (BCR) complex, comprised of antigen recognition and signaling components, functions in initiating B cell activation. While structural studies have described BCR domain organization, gaps remain in our understanding of its antigen binding domain (Fab, fragment antigen-binding) disposition, and how antigen binding is sensed to initiate signaling. Here, we report antigen affinity and signaling of the immunoglobulin (Ig) class IgM and IgG BCRs and define conformational states of full-length BCRs of two human broadly neutralizing antibodies, the glycan-specific, heavy chain domain-swapped, I-shaped 2G12, and a canonical Y-shaped antibody, CH31, that recognizes the CD4-binding site on the HIV-1 Envelope protein (Env). The BCRs adopted the shapes (I or Y) of their respective soluble antibodies, and both Ig class of BCRs of the same specificity bound Env trimers with similar affinities. We observed antigen-valency dependent differential signaling by the 2G12 IgM and IgG BCRs with trimeric Envs. Cryo-electron microscopy of the 2G12 IgG and CH31 IgM BCRs revealed varied Fab orientations. Structural comparisons revealed hinge points and regions of flexibility in the BCRs suggesting a highly dynamic structure of the BCR complex. Taken together, our results provide an integrated understanding of BCR structure, conformation, antigen recognition and signaling, and provide the basis for understanding antigen induced BCR signal transmission.

**One Sentence Summary:** Cryo-electron microscopy structures of full-length B cell antigen receptor complex provide a novel dynamic BCR model and a basis for understanding antigen binding induced signal transmission.

## INTRODUCTION

B cell activation is initiated by the binding of an antigen to the B cell antigen receptor (BCR) that is assembled and expressed on the B cell surface as a complex (BCR complex) with the disulfide-linked Igα/Igβ (CD79a/CD79b) ^1–3^. The specificity of the antigen binding domain (Fab, fragment antigen-binding) is formed by the combination of V_H_/V_L_ (Variable-Heavy/Variable-Light) sequences while the Igα/Igβ constitutes the signaling component with its conserved cytosolic ITAM (immunoreceptor tyrosine-based activation motif) tail that gets phosphorylated and initiates the signaling events leading to B cell activation^4,5^. Structural information of the BCR complex in its resting auto-inhibited state and its transition to the antigen-bound signaling-competent state are key to understanding BCR oligomerization and the mechanism of signal transduction during B cell activation ^2,6–8^. Cryo-electron microscopy (cryo-EM) structures of a human ^9,10^ and a mouse ^11^ BCR provided a general principle of the assembly of the BCR complex, particularly the organization of the extracellular domains (ECD) and the transmembrane domains (TMD) and the atomic details of the residue contacts that form the association of BCR with Igα/Igβ. Key features of the first BCR structures include the validation of the previous biochemical evidence of the asymmetry of BCR/ Igα/Igβ association ^2,12^, the interactions of the TMD domains of BCR Heavy chain (HC)/ Igα/Igβ to form a four-helix bundle ^9,10^ and the distinct modes of the ECD interactions in the IgM and IgG class BCRs^9^. These structures, however, provided limited information on the Fab domains, primarily due to domain flexibility and the associated poor EM density, and it remained unclear how antigen binding related conformational changes could be transmitted across the ECD and TM to initiate signaling. While the Fab flexibility was noted in each of the reported BCR structures, the conformational variability of the BCR structure was not studied and the reconstructed BCR structures present an overall rigid model of the BCR complex in its resting state and the role of BCR conformational state dynamics in signal transduction remains unresolved.

A category of HIV-1 broadly neutralizing antibodies (bnAbs) that bind to closely spaced glycans on the HIV Envelope (Env) protein adopt an I-shape Fab configuration ^13–18^ which is unlike the Y-shape of conventional antibodies. This unique structural category is called Fab-dimerized glycan (FDG) antibodies^17^. Among FDG antibodies, the side-by-side Fab dimerization is achieved through interfacial hydrophobic interactions^17^. The bnAb called 2G12 is unique among FDG antibodies in that the I-shaped dimer configuration involves V_H_ Fab domain swapping ^16,19–21^. The Fab dimerization of FDG antibodies is essential for their avid binding to the HIV Env oligomannose shield and their biological function^14,19,22^. While soluble FDG mAbs can adopt the I-shape, there is only indirect evidence that IgG BCRs with 2G12 specificity also adopt the same configuration on the B cell surface, since the non-domain swapped form of BCRs (either a non-domain exchanged mutant or the germline 2G12 BCR) showed no signaling in response to either an Env trimer or synthetic glycoconjugates^14,22,23^. To date, how the FDG BCRs are configured on the B cell surface, the structural organization and interactions of the different Ig/signaling domains, and how antigen affinity is sensed for BCR signaling have not been studied.

We developed human B cell lines expressing either IgG or IgM class BCRs to study *in situ* B cell signaling and antigen binding to extracted BCRs, and to perform cryo-EM structural analyses. We focused on the specificities of two distinct HIV-1 bnAbs, 2G12 or CH31; the latter being a CD4 binding-site bnAb ^24,25^ in the same class as VRC01^26,27^, the BCR structures of which were recently described^9^. CH31 and 2G12 respectively represent the canonical Y-shape and the non-canonical I-shaped Fab configuration. The BCRs with V_H_/V_L_ sequences of each specificity were expressed on Ramos B cells with native signaling and co-receptor molecules, which is unlike the recombinantly expressed BCR complex used in previous structural studies^9,11^. Here we report full-length cryo-EM structures of 2G12 IgG and CH31 IgM class BCRs wherein we define the variability in Fab position and identify hinge points of BCR flexibility to illustrate a dynamic model of the full-length BCR complex with implications in BCR allostery and signaling.

## RESULTS

### Purification and biochemical analyses of BCR complex

Ramos B cell lines (human B cell lymphoma line) were developed that expressed BCRs with specificity of either 2G12^16^ or CH31^24^ bnAbs, each of IgM or IgG class and with C-terminus tags (TwinStrep + His) to facilitate purification. The B cell lines showed cell surface expression of BCR heavy chains, the relevant kappa (κ) light chains, the Igα/Igβ chains, and co-receptors (**Fig. S1A**). The BCRs displayed functional signaling by cross-linking antibodies (**Fig. S1B-C).** Following detergent solubilization, the BCR complexes (BCR + Igα/Igβ) were purified through a combined approach involving tag-specific affinity purification and size exclusion chromatography (SEC) (**Fig. 1A**), as detailed in the methods section. The same methodology was employed for purification of each class (IgM, IgG) of either 2G12 or CH31 BCRs (**Fig. 1B**). The molecular sizes of the BCR subunits in the purified BCR fractions were confirmed by SDS-PAGE and each BCR complex components (H-chain, κ L-chain, Igα, Igβ) identified by western blot analysis (**Fig. 1C**), thus confirming the integrity of the BCR complex associated with the signaling component in the SEC eluted BCR fractions (**Fig. 1B**). In SPR binding assays, each of the detergent solubilized BCRs bound specifically to Env gp140 trimer in a dose-dependent manner (**Fig. 1D**). Env binding to 2G12 BCRs binding was weaker than to CH31 BCRs, which is consistent with the binding to the soluble antibody forms (**Fig. 1D**).

**Fig 1.**
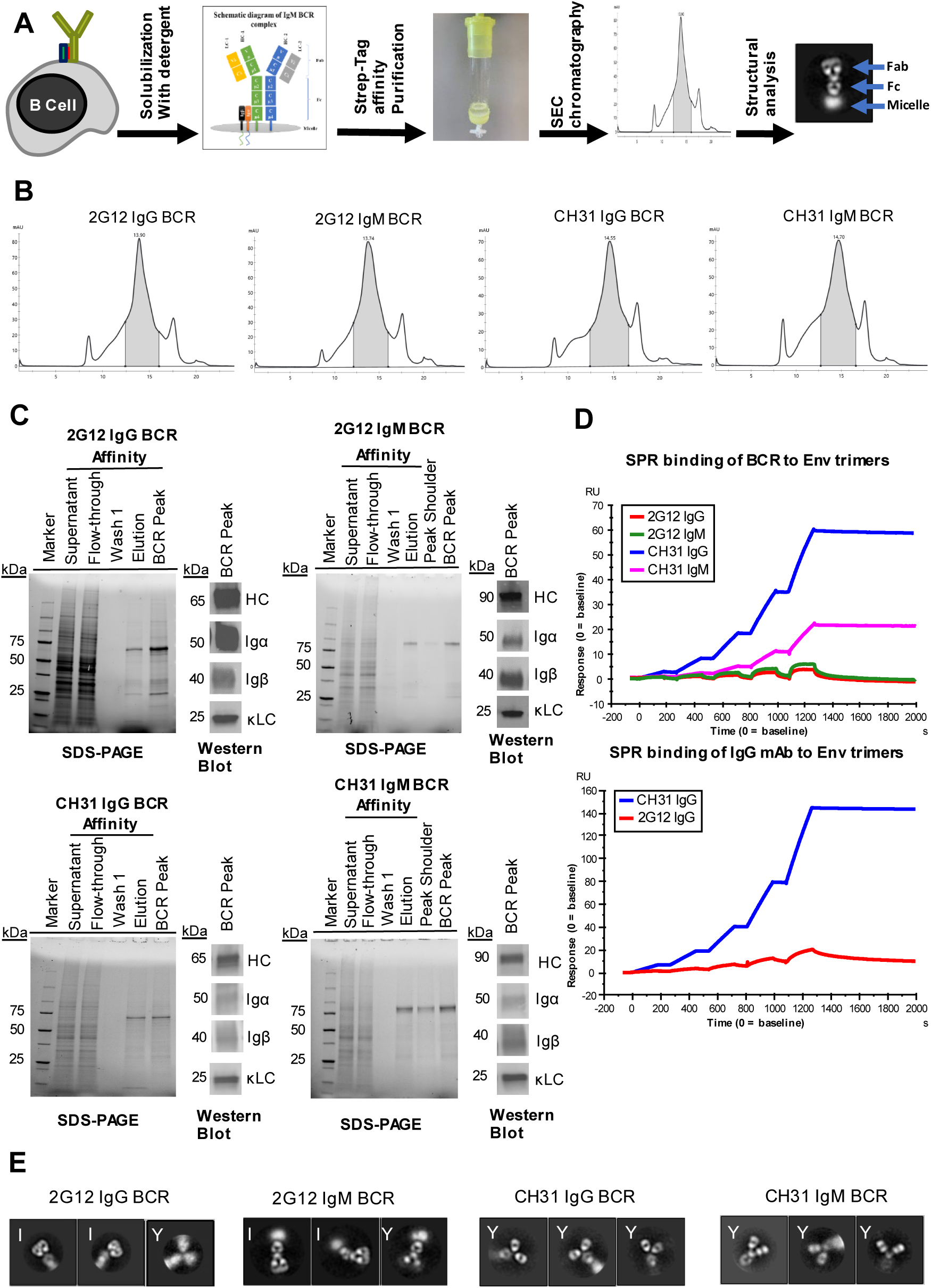
Expression, Purification and Biochemical analysis of B cell Receptor (BCR) from Ramos B cells. **(A)** Schematic of BCR expression and purification workflow. Following detergent solubilization of Ramos cells expressing either 2G12 or CH31 specific IgG or IgM class BCRs with a Strep-tag, BCR complex proteins were purified using twin-Strep-tag affinity purification and size exclusion chromatography (SEC) for biochemical and structural analysis. **(B)** SEC chromatography profile of BCR proteins following affinity purification. Shaded region in each plot shows the eluted fractions of the BCR complex. **(C)** SDS-PAGE data shows protein staining for each indicated fractions and Western blot analysis with BCR subunit specific antibodies reveals the immunoglobulin (heavy chain, HC), light chain (kappa light chain, kLC), and the associated signaling (Iga/Igb) subunits of the intact BCR complex. **(D)** Surface plasmon resonance (SPR) titration curves of purified BCR [top panel] and soluble IgG [bottom panel] of CH31 or 2G12 specificity to HIV Envelope (Env) trimer (CH505TF v4.1). Env trimer was injected over BCR or mAb immobilized sensor surface in a two-fold dilution series ranging 2 – 80 nM. **(E)** Negative stain electron microscopy (NSEM) analysis of IgG and IgM class BCR complexes. The NSEM class average images show representative BCR shape configurations (I or Y-shape) of 2G12 and CH31 specific BCRs.

Negative stain electron microscopy (NSEM) analysis showed that both IgM and IgG class of purified 2G12 BCRs embed in lipid micelle and adopt predominantly I-shaped Fab architecture, although in 2D class-averages a small proportion of Y-shaped BCR was also observed (**Fig. 1E, S2**). Thus, BCRs of the domain-swapped 2G12 specificity predominantly adopt the I-shape consistent with the shape configuration of soluble 2G12 IgG mAbs^13^. In contrast, both the IgG and IgM BCRs of CH31 were Y-shaped (**Fig. 1E, S2**), which is the conventional configuration of a soluble antibody. Thus, the purified BCR complexes were intact in a membrane associated form, retained their antigen binding specificity and adopted the shape configuration (I-or Y-shape) of the respective antibody counterparts.

### Purified BCR Affinity to Env trimers

We compared the binding of soluble HIV Env gp140 trimers to both 2G12 and CH31 BCRs to that of their soluble forms including Fabs and mAbs. We tested 5 different Env trimers that bound with differing affinities to 2G12 Fab dimer, ranging from 0.45μM to 0.9nM (**Fig. 2**). When comparing the two distinct BCR classes, each of the trimers bound to either 2G12 IgM or IgG BCRs with similar affinities and notably with the same hierarchy in trimer affinity discrimination. However, the kinetic rates (association and dissociation rates) were faster for both 2G12 BCR classes when compared to the soluble IgG mAb (**Fig. 2A-B, S3, Table S1**). For CH31, the selected trimers bound to both classes of CH31 BCR with similar affinities, but each with higher affinities than the binding of CH31 IgG mAb due to faster binding kinetics (**Fig. 2, S4, Table S2**). When compared to the binding of either CH31 Fab or mAb, the same hierarchy in BCR binding association rates was observed (**Fig. 2A, S4, Table S2**) and that is consistent with our previously reported finding that CH31 BCR signaling is dependent on trimer association rate^28^. In contrast, each of the Env trimers bound to 2G12 BCRs with similar and fast association rates (>10^5^ M^-1^s^-1^) (**Fig. 2A**). Overall, these data showed that the trimer affinity discrimination by BCRs followed the same trend as that of soluble antibodies, and both IgG and IgM class BCRs of the same specificity bind trimers with similar kinetic rates and affinities.

**Fig. 2.**
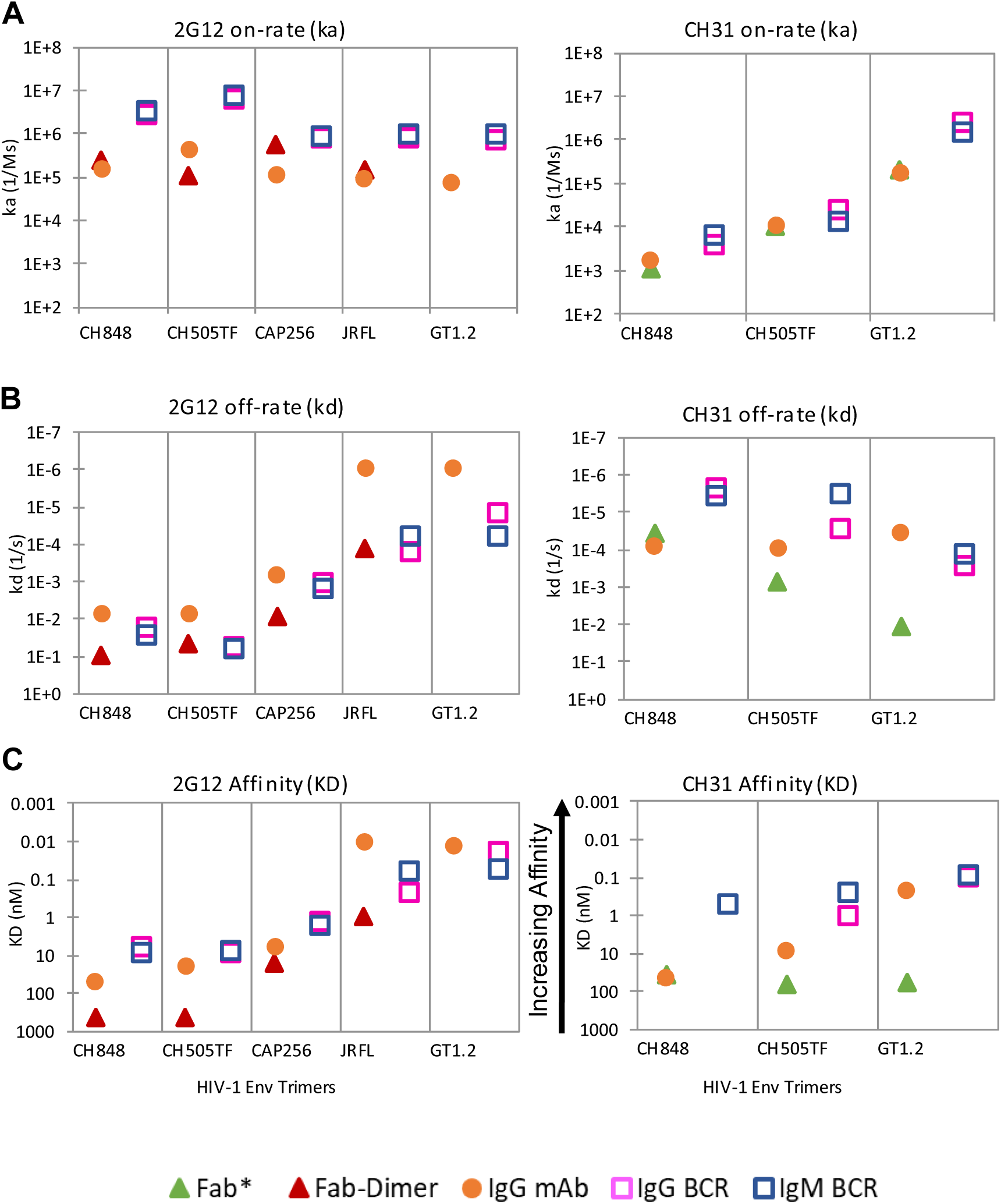
Affinity (K_D_) and Kinetic rates (k_a_, k_d_) of IgM and IgG BCRs to HIV-1 Env Trimers. **(A)** Association rates (k_a_) of 2G12 [left panel] and CH31 [right panel] to soluble HIV-1 Env gp140 trimer proteins. Measurements were done using Fab [green triangle], Fab-Dimer [red triangle], IgG mAb [orange circle], IgG BCR [pink square] and IgM BCR [blue square]. **(B)** Dissociation rates (k_d_) of 2G12 and CH31 to Env trimers. Data point labeling as described in A. **(C)** Dissociation constant (K_D_, affinity) of 2G12 and CH31. Data point labeling as described in A. Surface Plasmon Resonance (SPR) was used to conduct titrations with soluble Env trimers. Env trimers were immobilized to chip surface with either Fab or Fab-Dimer injected over the surface. mAb and BCR were captured to chip surface with trimers injected over the immobilized surface. Injections were done in a two-fold dilution series with concentration ranging for Fab 200 – 8000nM, Fab-dimer 12.5 – 400nM and Env trimer 2 – 1000nM. Curve fitting made use of either heterogenous ligand model or 1:1 Langmuir model with global R_max_. Faster kinetic parameters are reported in the case of heterogenous ligand model. Data points are the averages of at least three independent measurements. Data of 2G12 Fab-Dimer affinity to BG505 GT1.2 Env trimer are provided in supplemental data (Table S1). *For comparison, CH31 Fab kinetics and affinity data to CH505TF v4.1 and BG505 GT 1.2 Env trimers are plotted from previously published data^28^.

### Activation of B cells expressing 2G12 IgM or IgG BCRs

Previously we reported that B cells expressing IgM BCRs of CH31 sense antigen affinity based on association rates, rather than affinity (K_D_)^28^. Thus, we hypothesized that all the Env trimers tested that bound with similar association rates but different affinities (**Fig. 2**) to 2G12 BCRs would each strongly trigger 2G12 BCR signaling. We tested a panel of Env trimers that bound the 2G12 IgG ectodomain with differing affinities (K_D_ ranging from about 0.5 μM to 1nM) for calcium flux (Ca2+) responses, a measure of B cell activation, in Ramos cell lines expressing either 2G12 IgM or 2G12 IgG BCRs (**Fig. 3**). As we had predicted, each of the trimers that bound 2G12 BCRs with similar fast association rates (k_a_>10^5^ M^-1^s^-1^) (**Fig. 2A**) showed strong Ca2+ responses (average peak response >60% of control, peak response range 65.5 – 115.3%) in 2G12 IgM B cells (**Fig. 3A, Table S3A**). Although a trend showing relatively stronger signaling with higher trimer affinity was observed (**Fig. 3A**). The multimeric form of the trimers also induced strong Ca2+ responses in 2G12 IgM cells (**Fig. 3B**). In contrast, each of the same trimers induced about 2-3-fold weaker Ca2+ response (<50% of control, peak response range 26.4 – 48.3%) in the 2G12 IgG B cells (**Fig. 3C, Table S3A**). 2G12 IgG B cells, however, gave stronger Ca2+ responses (about 2-fold higher peak response) when stimulated with multimers of trimers (**Fig. 3D, Table S3B**). By contrast, the CH31 IgG B cells did not show enhancement in Ca2+ response to the multimer-of-trimer versions of the Env antigen relative to the trimeric version, and as reported previously^28^, gave a stronger Ca2+ response with trimers that bound with faster association rates (**Fig. S5**). Thus, while both trimeric and multimer-of-trimer forms of the antigens strongly activated 2G12 IgM B cells, the activation strength in 2G12 IgG B cells was dependent on the antigen valency, consistent with the recently reported class-specific antigen sensing by B cells^29^. In addition, since the IgM and IgG BCRs bound to Env trimers with nearly identical affinities and kinetic rates, we hypothesize that the BCR paratope-antigen binding interactions on B cell surface are not distinct for the two BCR classes and that the observed class-specific sensing of antigen has a structural basis^9^ that results in distinct signal transduction.

**Fig. 3.**
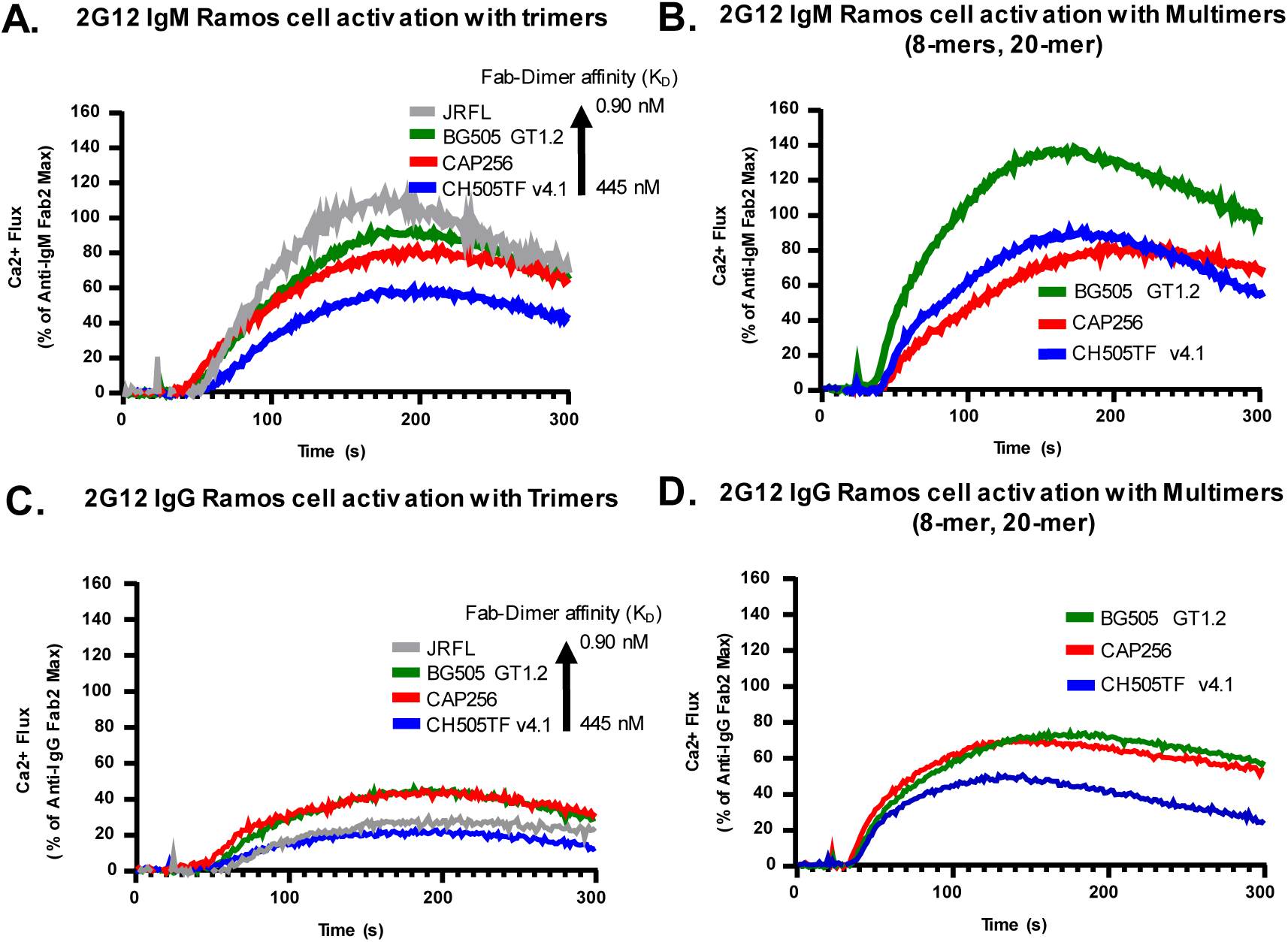
Activ ation of Ramos Cells expressing IgM or IgG BCRs with specificity of 2G12 or CH31. **(A)** Calcium (Ca2+) flux responses over time (s) induced in 2G12 IgM Ramos cells by Env trimer proteins. 2G12 IgM Ramos cells treated with either BG505 GT1.2 (green), JRFL (gray), CAP256 (red), or CH505TFv4.1 (blue) Env trimers at 30nM and monitored for intracellular Ca2+ binding dye fluorescence for 300 s. **(B)** Ca2+ flux responses in 2G12 IgM Ramos cells activated by Env trimer multimers including BG505 GT 1.2 20-mer (green), CAP256 8-mer (red), CH505TFv4.1 8-mer (blue) at concentrations of 30nM. Env trimers and their multimeric forms prepared at the same per unit trimer concentrations for B cell activation (30nM). Ca2+ flux results for the 2G12 IgM Ramos cells are presented as a percentage of the maximum anti-human IgM F(ab’)_2_ response and are an average of at least two measurements. **(C)** Ca2+ flux responses induced in 2G12 IgG Ramos cells by Env trimer proteins including BG505 GT 1.2 (green), JRFL (gray), CAP256 (red), CH505TFv4.1 (blue) at 30nM. **(D)** Ca2+ flux responses in 2G12 IgG Ramos cells by Env trimer multimers. Overlaid responses shown are with BG505 GT 1.2 20-mer (green), CAP256 8-mer (red) CH505TFv4.1 8-mer (blue) at 30nM. 2G12 IgG Ramos cells were treated with SOSIP trimers or multimers identically to the 2G12 IgM Ramos cells. Ca2+ flux results for the 2G12 IgG Ramos cells are presented as a percentage of the maximum anti-human IgG F(ab’)_2_ response and are an average of at least three measurements.

### Cryo-EM structure of 2G12 IgG BCR

To visualize the architecture of the 2G12 BCR we performed single particle cryo-EM analysis on 2G12 IgG BCR that were detergent extracted and purified from Ramos B cells **(Fig. 4, S6-S7, Table S4**). Representative 2D classes revealed evidence suggesting considerable conformational heterogeneity (**Fig. 4A**). In a smaller subset (∼10%) of particles we could visualize the entire BCR molecule with features that could be unambiguously assigned to the domain-swapped 2G12 Fab, the Fc and the signaling components Igα and Igß. The micelle appeared as an indistinct oval at the position where the transmembrane helices would be located. The hinge region between the Fab and the Fc was also visible, albeit faintly, in the 2D classes (**Fig. 4A**) and in the ab initio reconstructions (**Fig. S6 I and J**). This region appeared highly flexible, resulting in different orientations of the 2G12 Fab relative to the Fc (**Fig. 4B-E**). In addition to the complete view of the 2G12 BCR in a small subset of 2D classes, there were several views among the 2D classes where either the Fab and the Fc were visible, or the Fc, Igα/β and micelle were visible. This is consistent with the hinge flexibility mentioned above, that resulted in diverse conformations of the 2G12 BCR in the specimen, with the refinements aligning on different regions of the BCR depending on the relative orientation of its components.

**Fig. 4.**
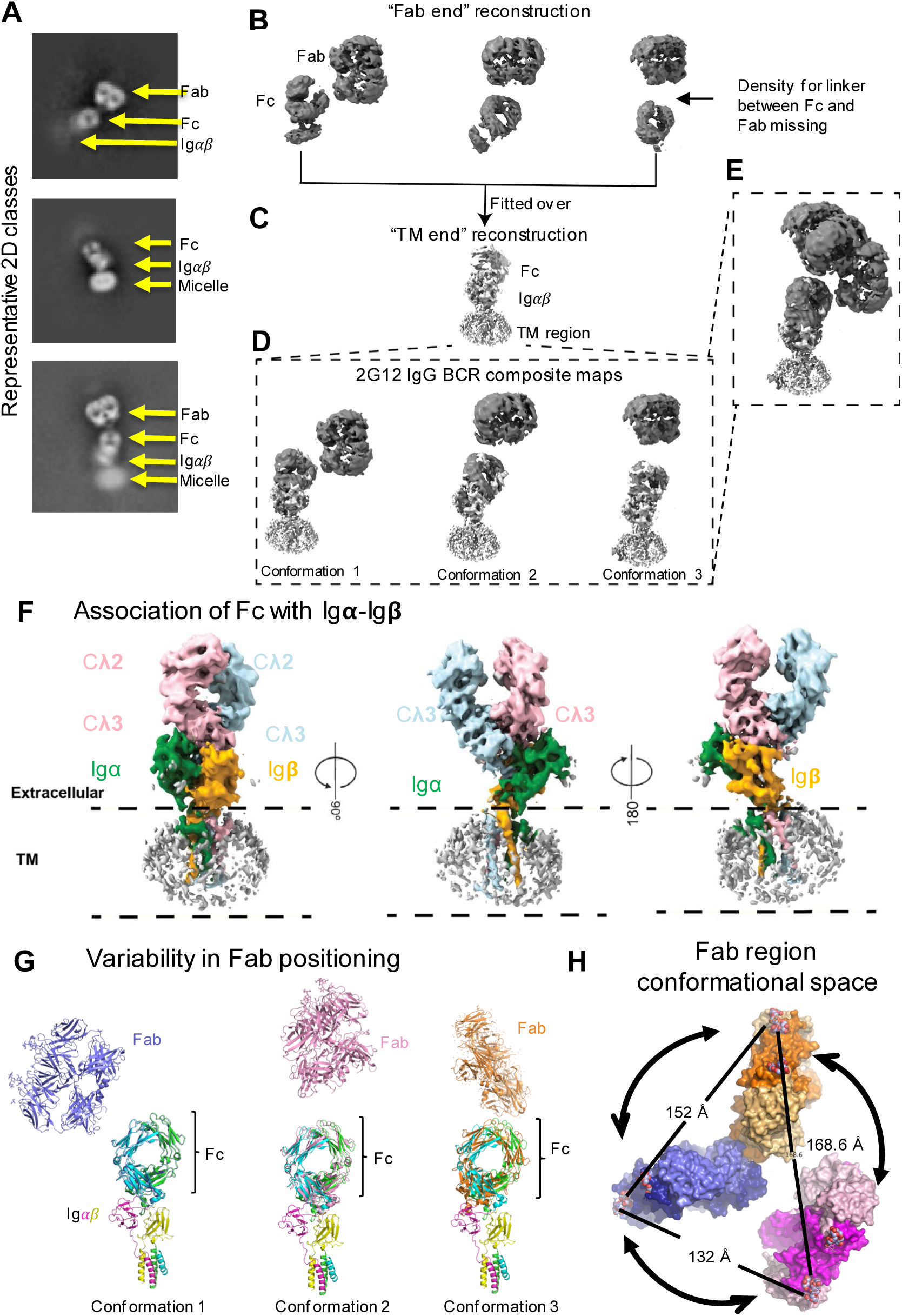
Cryo-EM structure of 2G12 IgG BCR. **(A)** Representative 2D class averages with different regions of the BCR labeled. **(B)** Three distinct populations from focused refinement and classification of the “Fab end”. **(C)** Focused refinement of the “TM-end” of the BCR. **D.** Composite reconstructions of the 2G12 BCR obtained by combining the “Fab end” reconstructions with the “TM-end” reconstruction. **(E)** Superposition of the volumes shown in panel **D**. **(F)** The “TM end” reconstruction colored by components, with the Fc chains colored pink and cyan, Igα colored green and Igβ colored orange. **(G)** Coordinate fits to the composite maps. **(H)** The structures shown in panel **G**. overlaid by the Fc and viewed top down from the “Fab end”. The heavy chains are colored magenta, dark blue and orange, and the light chains are colored pink, light blue and light orange. A crystal structure of glycan-bound 2G12 Fab was used for map fitting. The bound glycans are shown as spheres. Distances were measured between the distal glycan binding sites to obtain an estimate of the range of Fab motion on the 2G12 IgG BCR.

The region encompassing the Fc, Igα/β and the micelle enclosed transmembrane regions was refined to a resolution of 4.5 Å, with the Fc and Igα/β ectodomain further locally refined to a resolution of 4.4 Å (**Fig. S7**). Similar to the published BCR structures, we observed an asymmetric structure with one Igα/β pair associated with the Fc (**Fig. 4F**). The published structures of the IgM and IgG BCRs differed markedly in their Fc ectodomain configurations, with a 10-residue Igα loop spanning residues V69 and E79 driving the differences (**Fig. S8**)^9^. In the IgM BCR, this loop stacked against and contacted one of the two Fc chains, while in the IgG BCR the loop inserted between the two chains. This dramatic shift in the loop position impacted the conformation of other regions on Igα including its TM region, which was shifted by ∼6 Å, measured at residue R166, causing a tilt in the Igβ TM. As expected, our 2G12 IgG BCR structure resembled the published IgG BCR structure in its relative disposition of the Fc with respect to the signaling components. Despite the overall resemblance, conformational differences were noted, particularly in loop regions, including the key 69-79 Igα loop that contacts the Fc, the loops connecting the ectodomain to the TM regions, as well as the hinge region in the Fc (**Fig. S9**).

Unlike in the previously published structures, here we defined the Fab regions of the 2G12 IgG BCR. The 3D reconstructions of the 2G12 BCR revealed I-shaped configuration of the 2G12 Fab arms, with flexibility in the linker region between the Cg1-Cg2 domains resulting in the movement of the Fab region while the individual Fab moieties remained stably associated with each other in their I-shaped configuration. We obtained three reconstructions of the Fab region along with the Fc, and in some cases with portions of density consistent with Igα/ß associated with the Fc regions (**Fig. 4B** and **S6J**). Since the Fc appeared as a common motif between the “Fab-end” and the “TM-end” reconstructions, we aligned the structures by the Fc region to obtain three distinct composite maps of the complete 2G12 BCR (**Fig. 4C-E**). The three reconstructions differed in their orientation of the Fab relative to the rest of the BCR and spanned ∼130-170 Å separation between the Fab dimers in the different conformations (**Fig. 4G-H, Movie S1**). Although, the extent of BCR Fab domain motion may be impacted by the membrane mimetic chosen for this study, these structures establish the connection between the Fab and the Fc as a major hinge region about which the Fab can sample substantial variations in its orientation. The asymmetric configuration that arises from the interaction of a single Igα/ß pair with one of the two Fc chains also introduced anisotropy in the range of motion that the 2G12 Fab could sample about the Fc, with the Fab orientation tilted away from the Igα/ß side of the BCR.

Thus, our results provide the first views of a complete BCR structure, including details of the conformational diversity of the Fab region. The dynamic nature of the BCR suggested by our structural analysis are relevant to its antigen searching and sensing properties. The regions of flexibility identified in the BCR structure, centered around a malleable Igα loop, provide a path for the antigen binding signal to transmit from the Fab to the ITAM.

### Cryo-EM structure of CH31 IgM BCR

We next determined the structure of the CH31 IgM BCR by single particle cryo-EM analysis (**Fig. 5 and S10**). Unlike the unique domain-swapped I-shaped configuration of 2G12, the CH31 antibody adopts a canonical Y-shaped configuration, both in the soluble ectodomain version of the IgG as well as in the BCR (**Fig. 1E**). The 3D reconstruction of full length CH31 IgM BCR revealed well-defined densities for the Fab regions that confirmed a Y shaped configuration of the immunoglobulin ectodomain. Considerable flexibility was observed in the linker region of Cμ1-Cμ2, resulting in the two Fab arms adopting different conformations with different distances between them (**Fig. 5A-E**). Along with this region of flexibility, the linker region between Cμ2-Cμ3 added another dimension of flexibility resulting in the movement of the entire Fab region. We followed a similar strategy as we have done for the 2G12 BCR, of separate focused refinement of the “Fab-end” and the “TM-end” of the BCR, followed by alignment of the Fc regions that were common between the two to obtain 3D reconstructions of the full-length BCR (**Figs. 5 B-E**). We observed considerable diversity in the orientation of the Fab arms, with suggesting flexibility in the connection between the Fab and the Fc that resulted in the Y-shaped Fab region “swaying” about the Fc. We also observed different relative orientations of the two Fabs, which is distinct from the 2G12 BCR, where the rigid domain-swapped configuration of the Fab precluded this additional diversity in the BCR conformational ensemble.

**Fig. 5.**
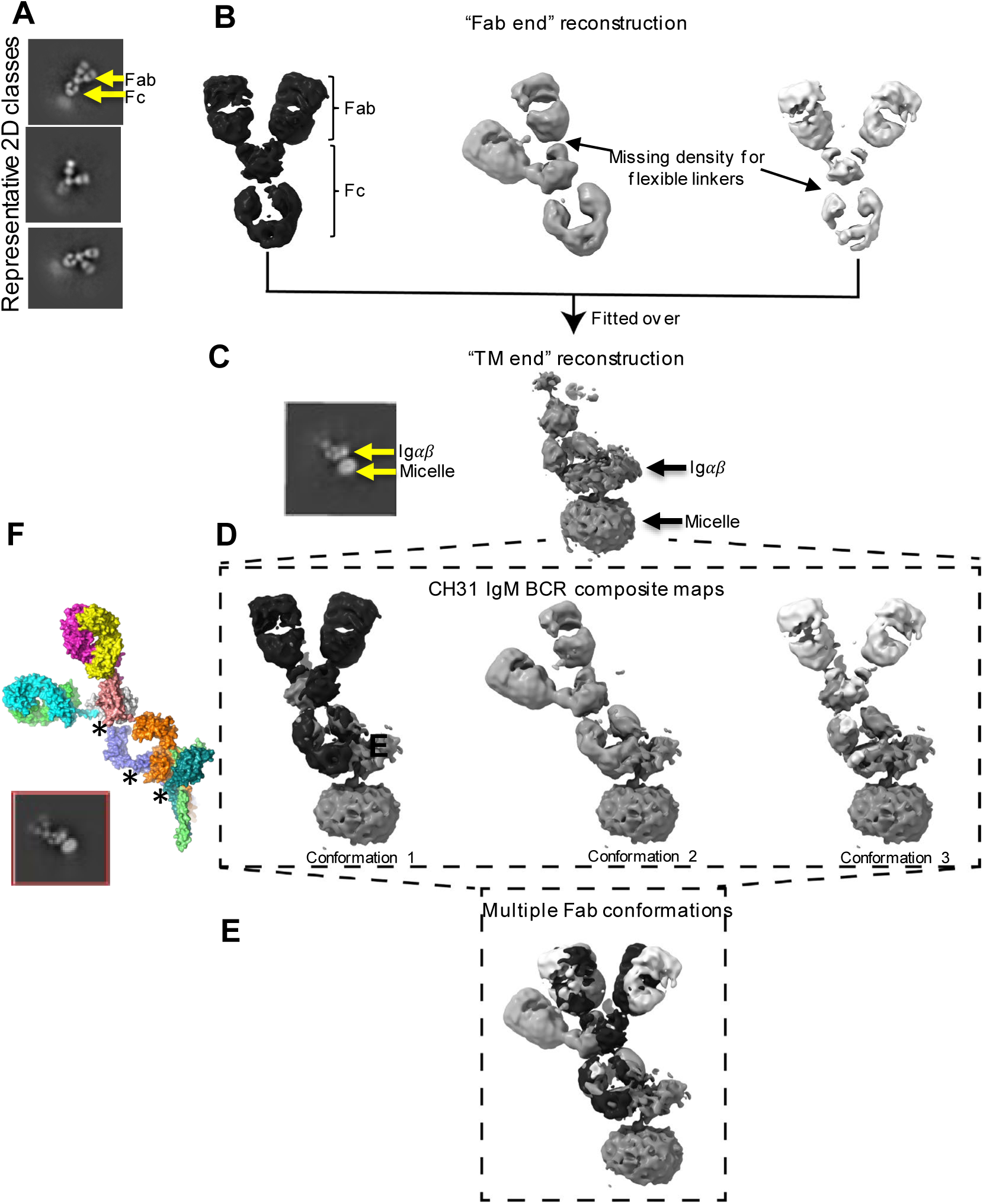
Cryo-EM structure of CH31 IgM BCR. **(A)** Representative 2D class averages showing variability in Fab orientations. **(B)** Three distinct populations from focused refinement and classification of the “Fab end”. **(C)** Focused refinement of the “TM-end” of the BCR. Inset is a 2D class showing a view of the “TM end”. **(D)** Composite reconstructions of the 2G12 BCR obtained by combining the “Fab end” reconstructions with the “TM-end” reconstruction. **(E)** Superposition of the volumes shown in panel **D**. **(F)** Fitted model of CH31 IgM BCR shown in surface representation. The asterisks indicate hinges about which domain motions occur in the BCR. Inset is a 2D class showing a view of the full CH31 IgM BCR.

Our recent molecular dynamics (MD) simulation study on the CH31 BCR supports a dynamic model of the membrane associated BCR and that antigen binding can allosterically induce changes in BCR conformations, including TMD helices^34^. The antigen binding induced conformational changes proposed in that MD study are compatible with our cryo-EM BCR dynamic model. MD of the same BCR complex without an antigen, when carried out in a POPC (1-palmitoyl-2-oleoyl-sn-glycero-3-phosphocholine) bilayer, are consistent with the motions observed in the cryo-EM structures. As shown in **Fig. S12 (Movie S2),** Fab arms are mobile and TMDs exhibit dynamical motions.

Taken together the structures of the 2G12 IgG and CH31 IgM BCRs show that a dynamic Fab region is a hallmark of BCR structures, irrespective of the BCR class.

## DISCUSSION

Our studies provide an integrated understanding of BCR antigen binding, BCR signaling and cryo-EM structures of the BCR complex with specificity of two key HIV-1 bnAbs that adopt distinct Fab shape configuration. While further validating the general assembly of the asymmetric BCR structure as described previously^9–11^, our full-length structures of two different BCR class (IgM, IgG) together provide a distinct and dynamic model of the BCR complex. Specifically, we have visualized different BCR conformational states and defined multiple hinge points centered around the Igα loop that affords flexibility to both the TMD and the ectodomains, and thereby present a flexible BCR structure with the potential to change upon antigen binding. Importantly, our dynamic BCR model provides a structural basis for understanding the transmission path of the conformational signal from the BCR ectodomains to the cytosolic ITAM.

In the previously described human as well as mouse BCR structures^9–11^, the Fab dynamics were noted but not studied primarily due to poor resolution of the Fab domains. Therefore, these structures provided an overall ‘rigid’ model of the BCR complex that is likely sampling a single conformational state of a highly flexible BCR complex. Here we describe the first cryo-EM visualization of the flexibility of the BCR complex, and map multiple conformations in the CH1-CH2 as well as within Cμ3-Cμ4 (in CH31 IgM) and that together likely allow the hinge and the loop regions to transmit conformational changes from the ectodomains to the TMD of the BCR complex. The identification of multiple hinge points in the BCR structure is consistent with the reported flexibility of soluble antibody domains that include antibody hinge region between the Fab and F_c_, as well as the relative disposition of the Fab to the constant domains^30–33^. Overall, the structural comparisons indicate a dynamic and flexible BCR structure where key hinge and loop regions can orchestrate the relay of conformational changes between distant regions of the structure. A central role is played by the Igα component that demonstrates remarkable ability to adapt its interactions to different BCR classes and its malleable nature makes the Igα a likely central player in transmitting antigen binding related conformational signals to the TM and the ITAM motifs.

Based on our BCR model, the hypothesized changes in BCR flexibility can lead to either disruption of BCR self-association^2,35^ or association with a regulatory co-receptor molecule^36^ and subsequently transition the BCR from a ‘resting’ to an ‘activated state’. Notably, the proposed autoinhibition potentially involving an ITAM motif fold-back onto Igβ TMD^11^ can be allosterically exposed for phosphorylation following antigen binding. Thus, our dynamic BCR model supports a conformation-induced activation model and provides the structural basis for understanding signal transmission. As in previous BCR structures, a limitation of our model is the presentation of the BCR complex in a detergent micelle. Future studies will further elucidate the role of membrane lipid compositions in the orientation, dynamics, and signaling of the BCR complex.

In this study, we report BCR structures of two distinct categories, one that adopts the conventional Y-shape (CH31) and a second that utilizes Fab domain swapping for an I-shape configuration (2G12). Previously, it was speculated that 2G12 BCR on the B cell surface could adopt the I-shape since its binding to viral Env epitope and neutralization function was dependent on the acquisition of the dimerization motif ^22,23^. In this study we provide direct evidence that like the soluble antibody form, the 2G12 BCRs purified from cell surface also adopt the Fab I-shape configuration and therefore utilize both Fab arms to avidly bind to closely spaced glycans. The observation that 2G12 BCRs could be strongly activated by the trimeric antigen forms that bound with different affinities but each with fast association kinetics, further validates our previous reporting on the role of association kinetics on B cell activation ^28^. While the domain-swapped Fab in 2G12 BCRs form a single paratope, bi-valent antigen binding requirement is not restricted to FDG specificity, since BCRs that adopt the canonical Y-shape, like CH31, are not activated by monomeric antigens, including antigens that bind with high affinity^28^. Thus, a minimal antigen binding stoichiometry of n=2 is required for B cell activation^28,37^ and both Fab arms of a conventional Y-shaped BCR likely need to be bound to the activating antigen^34^, either on different protomers of a trimeric antigen or on multimers of a monomeric antigen^28^. The above stoichiometry requirement is not inconsistent with the observation that the relative orientations of the Fabs in the CH31 BCR structures allowed far more conformational diversity than the more rigid domain-swapped orientation of 2G12 that utilizes both Fabs to engage a closely spaced glycan epitope. Thus, the greater mobility of CH31 Fabs likely favor scanning and engagement of epitopes on the antigen molecule by each Fab domain. It follows from above, that with the imposed entropic hurdle in engaging a highly dynamic Fab domain, antigens that bind with fast association kinetics will be favored for BCR binding and signaling, as we observed here and in our earlier studies^28^.

We have measured direct binding of purified BCRs to relevant HIV Env trimers and report affinity and kinetics data not previously studied. In addition, since the IgM and IgG BCRs bound to Env trimers with nearly identical affinities and kinetic rates, we hypothesize that the BCR paratope-antigen binding interactions on the B cell surface are not distinct for the two BCR classes and that the observed class-specific sensing of antigen^29^ has a structural basis that likely results in distinct signal transduction. Recently, we reported that the CH1 domain of IgG has a dominant role in both BCR expression and sensing of antigen valency^29^ and that was consistent with the observation that 2G12 IgG expressing B cells gave stronger activation with multimeric than with trimeric forms of antigen. As in the published BCR IgM and IgG class structure^9^, we observed Fc ectodomain configurations that were distinct. In the 2G12 BCR structure, we further note key conformational flexibility in multiple loops that include those in Igα, the TMD, as well as in the Fc, and therefore, a potential path for signal transmission.

Finally, our use of a B cell line to purify BCRs avoid the need to recombinantly express either the signaling component or the co-receptors that interact with the BCR complex on a non-B cell surface. Since BCRs are present on protein islands that are associated with distinct coreceptors in the resting and activated state^38^, our cryo-EM analysis was performed on BCRs that were expressed on cell membrane with the lipid composition and associated with co-receptors that are native to a resting B cell surface. Our current studies to purify BCRs in antigen bound forms and from antigen activated B cells will provide a clearer understanding of the mechanism of signal transmission based on our dynamic BCR structure model.

## MATERIALS AND METHODS

### Construct design and stable cell line development

The Ramos cell line (RA 1 and cataloged as ATCC CRL-1596) were modified through stable transfection to express specific immunoglobulin B cell receptors (BCR), comprising both heavy and light chains, as outlined in previous studies^25,28,39^. Briefly, heavy and light chains encoding genes are cloned separately in FEEKW expressing vector^48^ using Xba1 and BamH1 restriction sites.

In our study, we engineered a heavy chain construct by adding a Twin-Strep + His tag at the C-terminus. This modification was designed to facilitate the affinity purification process, enabling the isolation of BCR complex from the cellular membrane while maintaining its functional integrity. To validate the effectiveness of this modification, we conducted thorough QC analyses of expression, signaling and activation analyses. These experiments confirmed that the introduction of the Twin-Strep + His tag did not interfere with the normal functioning of the BCR for subsequent studies. Recombinant Ramos cells, 2G12 bNAb and CH31 specific IgM and IgG BCR expressing cell lines, served as the foundation for our in-depth exploration into various cellular processes, including cell signaling and activation mechanisms. Additionally, the modified BCRs with the attached strep-His tag provided a robust platform for BCR purifications, enabling us to isolate and analyze the receptor.

### Surface Expression and Functionality

After stable transfection of naive Ramos cells using lentivirus, cells were stained with fluorescent antibodies targeting the Fcɣ and ĸ light chain portions of the BCR and phenotyped using flow cytometry. The cells were then enriched at least two times to select for the highest double positive expressers of BCR. A final phenotype was done to ensure expression was at the same level as other BCR expressing Ramos cell lines.

QC analysis was conducted on the final cells to ensure proper formation and functionality of the B-cell. Proper formation was determined by staining the cells using fluorescent antibodies targeting CD19, CD20, CD45, and CD79b and phenotyped using flow cytometry. Expression levels were compared to unstained cells. Proper functionality was determined by tracking the signaling components within the B-cells. Cells were incubated with anti-IgM or anti-IgG antibodies and fixed at 0, 0.5, 1, 2, 5, 10, 15, and 30 minute time points. The cells were then stained with antibodies targeting pSyk, pBtk, pBLNK, and pErk markers. Levels of these signals were then collected using flow cytometry. All cell lines referenced in this paper went through these QC experiments and all showed proper formation and function.

### Cell culture and harvesting

Ramos cells carrying a specific BCR were grown and collected using a standardized method. Initially, 10 million Ramos cells were placed in a T75 culture flask with 40 ml of 1X RPMI medium supplemented with 10% Fetal Bovine Serum (FBS). These cells were grown for three days at 37°C in a CO2 incubator until confluent. Then the cells were collected, and 50 million cells were transferred to a larger T850 culture flask containing 450 ml of 1X RPMI medium with 10% FBS. These were then incubated for an additional three to four days at 37°C.

After this second phase of cultivation, the cells were spun at 3000g for 20 minutes at 4°C in 500 ml centrifuge tubes to form a cell pellet. The supernatant was removed and the cells were rinsed with 15 ml of 1X PBS by gently resuspending the pellet. Then, the cells underwent another centrifugation at 2000g for 10 minutes at 4°C using 50 ml Falcon tubes to get a dense pellet. The PBS was removed without disturbing the pellet and was stored at −20°C until further processing.

### Purification of BCR complex

The purification process started by thawing the frozen cell (3-4 billion cells) pellet on ice and suspending it in 20 ml of Buffer A (25 mM HEPES, 150 mM NaCl, pH 7.5). After centrifugation at 2000g for 10 minutes at 4°C, the supernatant was removed, and the pellet was re-suspended in 10 ml of Buffer A. The mixture underwent homogenization in a Dounce homogenizer, followed by the addition of 10 ml of Buffer B (25 mM HEPES, 150 mM NaCl, 2% DDM, 1 protease inhibitor tablet, pH 7.5) for the final homogenization. This lysate was transferred to a 50 ml Falcon tube, left on a rocker at 4°C for 3 hours, and then centrifuged at 21000 g for 1 hour at 4°C. The resulting supernatant was collected, pooled, and filtered through a 0.45µm syringe filter before being stored on ice for further use.

A gravity flow column packed with 5 ml of Strep-Tactin (IBA) resin underwent two washes with Buffer A, followed by one wash with 10 ml of Buffer C (25 mM HEPES, 150 mM NaCl, 0.02% GDN, pH 7.5) and an additional 10 ml Buffer C incubation. After discarding Buffer C, the cleared lysate supernatant was added to the column and allowed to bind to the resin on a rocker at 4°C for 2 hours.

The column was then attached to a stand, allowing the supernatant to flow through for collection, indicating binding efficiency. The resin went through two washes with 10 ml of Buffer C and two washes with 10 ml of Buffer D (25 mM HEPES, 300 mM NaCl, 0.02% GDN, pH 7.5). BCRs were eluted from the resin using 20 ml of Buffer E (25 mM HEPES, 150 mM NaCl, 0.02% GDN, 4mM desthiobiotin, pH 7.5). The eluted sample was concentrated to 500 µl using a 100kDa MWCO Ultra centrifugal filter (Amicon) for subsequent SEC chromatography.

SEC purification utilized a Superose 6 Increase (10/300 GL) SEC column (Cytiva) with Buffer F (25 mM HEPES, 150 mM NaCl, 0.01% GDN, pH 7.5) at a flow rate of 0.5 ml/minute via an Äkta Pure system. Fractions corresponding to the BCR peak and shoulder peak were collected, pooled, and concentrated using a 100kDa cutoff to achieve a final protein concentration ranging between 1-3.5 mg/ml. Further characterization by NSEM showed that the primary BCR peak contained the highest quality complex.

Stability assessments were conducted on the purified BCR samples, revealing that the 14-day storage of samples at 4°C exhibited the highest homogeneity, with 64% of particles covered by class averages, rendering the remaining particles undetectable. Contrastingly, the fresh and 21-day-old samples demonstrated 38% and 50% homogeneity, respectively. Additionally, freeze-thaw experiments indicated a gradual decrease in homogeneity, with 45% and 28% homogeneity observed in the 1X and 5X freeze-thaw samples, respectively.

### SDS-PAGE and western blot analyses

This study investigated BCR components through SDS-PAGE and Western blot techniques to determine their composition and purity. Initially, samples from affinity purification and SEC chromatography fractions were concentrated using a 100kDa MWCO cutoff. Next, 6 µg of the concentrated protein underwent separation on stain free SDS-PAGE (BioRad) under reducing conditions, accompanied by unstained and stained protein ladders. The protein sample was mixed with 4X Laemmli sample buffer containing β-mercaptoethanol and boiled at 95°C for 10 minutes.

The separated proteins underwent gel imaging using ChemiDoc MP (BioRad), following which they were transferred onto a membrane using the Trans Blot Turbo Transfer system (BioRad). Post-transfer, the membrane was washed twice with 1XTBS buffer for 5 minutes and then blocked with 0.2% non-fat milk in 1XTBS for 1 hour at room temperature.

After blocking, the membrane underwent two washes with 1XTTBS buffer (0.1% Tween 20 in 1XTBS) and was incubated with primary antibodies diluted in Ab buffer (0.2% non-fat milk in 1XTTBS). Detection of IgM utilized goat anti-human IgM (biotinylated) at a 1:3000 dilution, while Kappa Light Chain employed goat anti-human kLC (biotinylated) at the same dilution. Igα detection utilized purified mouse anti-Igα at a 1:3000 dilution, and Igβ detection used purified mouse anti-Igβ at the same dilution. The membrane was sealed and incubated overnight at 4°C on a rocker.

Following primary antibody incubation, the membrane underwent three washes with 1XTTBS buffer for 5 minutes each. Subsequently, it was incubated with secondary antibodies prepared in Ab buffer. Biotin antibodies were detected using Avidin-AP at a 1:3000 dilution, mouse antibodies with Anti-mouse IgG-AP at the same dilution. The membrane was incubated for 1 hour at room temperature on a rocker.

After the secondary antibody incubation, the membrane was washed with 1XTTBS buffer for 5 minutes each. Excess buffer was removed, and an appropriate amount of Immuno-Star AP substrate was added to cover the protein site. Following a 5-minute incubation, excess substrate was removed, and images were captured using the ChemiDoc MP imager (BioRad) and analyzed through Image Lab software.

### Negative staining electron microscopy (NSEM)

A fresh aliquot from 4°C was diluted to 50 µg/ml with 0.02 g/dl Ruthenium Red in HBS (20 mM HEPES, 150 mM NaCl pH 7.4) buffer. After 5 min incubation, sample was applied to a glow-discharged carbon-coated EM grid for 8-10 second, blotted, consecutively rinsed with 2 drops of 1/20X HBS, and stained with 2 g/dL uranyl formate for 1 min, blotted and air-dried. Grids were examined on a Philips EM420 electron microscope operating at 120 kV and nominal magnification of 49,000x, and 34 images were collected on a 76 Mpix CCD camera at 2.4 Å/pixel. Images were analyzed by 2D class averages using standard protocols with Relion 3.0 ^40^.

### Fab and Fab-Dimer preparation from purified IgG samples

CH31 Fab and 2G12 Fab-Dimer were prepared using a modified protocol previously described ^47^. Purified IgG samples were dialyzed into 20mM sodium phosphate, 10m M EDTA, pH 7.0 and then concentrated to 20mg/ml using centrifugal filtration (Amicon). IgGs were digested using papain-agarose resin (Thermo Fisher Scientific) in 20m M sodium phosphate, 10mM EDTA, 20mM cysteine, pH7.4 for 5 hours at 37°C. Fab fragments were separated from undigested IgG and Fc fragments through a 1h incubation with rProtein A Sepharose Fast Flow (Cytiva). Finally, cysteine was removed by centrifugal filtration (Amicon) and buffer exchange to 1x PBS, pH 7.4.

Quality of Fab and Fab-Dimer was measured through SDS-PAGE and size exclusion chromatography (SEC). SDS-PAGE was conducted by loading 2μg protein/lane on a 4% to 15% TGX stain-free gel (Bio-Rad) nonreducing and 100 mM DTT reducing conditions. Loaded gels were run at 200V in Tris-glycine-SDS buffer. Bands were visualized using Chemidoc MP Imaging System or Gel Doc EZ imager (Bio-Rad), and band size was assessed with a protein standard ladder (Bio-Rad). SEC was performed by loading 15μg protein on to a Superdex 200 increase 10/300 column and run at 0.5ml/min using an Äkta Pure system (Cytiva). Fab peaks were analyzed with the Unicorn 7.6.0 software. Molecular weight was estimated using a linear regression calculated by running a mix of known-molecular-weight protein standards (Cytiva) on the same column.

### SPR binding studies

Affinity was measured through SPR using a BIAcore S200 instrument or T200 instrument (Cytiva) in HBS-EP+ 1x running buffer at 25°C. Two chip surface orientations were used to conduct measurements. The first orientation was used for the titration of Fab fragments to Env trimers. Biotinylated Env trimers were diluted to 1 – 2 μg/ml and injected at 5 μL/min over either a streptavidin immobilized CM5 chip or SA chip. 150 – 250 response units (RU) of ligand was captured on flow cells 2 – 4. The second orientation was used for the titration Env trimers to mAb and purified BCR. mAb and BCR were diluted to 3-50 μg/ml and injected at 10 μL/min over either anti-human Fc antibody immobilized CM5 chip for mAb samples or a SA chip for BCR samples. Ligand capture levels for mAb ranged from 130 – 380 RU and BCR capture levels ranged from 60 – 220 RU. Flow cell 1 was used as a blank surface reference for subtraction of nonspecific binding to chip surface, streptavidin, and anti-human Fc antibody. Ligand capture was followed by dissociation period of 30 mins or until response stabilized.

CH31 Fab was diluted to concentrations ranging between 200 nM and 8000 nM, while 2G12 Fab-Dimer was diluted to 12.5 – 400 nM. The following non-biotinylated Env trimers were used at the listed concentration ranges: CH505TF (2 – 200 nM), CH848 (10 – 1000 nM), GT1.2 (2 – 48 nM), CAP256 (1.5 – 48 nM), JRFL (2 – 48 nM). Analytes were injected at a flow rate of 50 μL/min over flow cells 1 – 4 using single-cycle kinetics injection type. Titration cycle consisted of five 180 s injections of Fab fragments at 2-fold increasing concentrations with 720 s dissociation after final injection. Chip surfaces were regenerated with Glycine, pH 1.5 or pH 2.0 for 30 s between cycles. HBS-EP+ running buffer titration cycle was used to account for signal drift in addition to the blank flow cell for double reference subtraction. Curve fitting of results was performed through BIAcore S200 evaluation software (Cytiva). Fitting made use of either heterogenous ligand model or 1:1 Langmuir model with global R_max_. Titration curves are representative of at least 3 data sets.

### Calcium Flux Measurements

Calcium flux experiments were performed using the FlexStation 3 Microplate Reader (Molecular Devices) in conjunction with the FLIPR Calcium 6 dye (Molecular Devices) as previously described^25^. A cell count for the Ramos cells was performed with a Guava Muse Cell Analyzer (Luminex) to ensure cell viability was greater than 95% and to calculate the volume of cells needed for a concentration of 1×10^6^cells/ml on the day of the experiments. The required volume of cells was then pelleted at 1500rpm for 5 minutes. The supernatant was then decanted and the cells were resuspended at a 2:1 ratio of RPMI media (Gibco) + FLIPR Calcium 6 dye (Molecular Devices). The cells were plated in a clear, U-bottom 96-well tissue culture plate (Costar) and incubated at 37°C, 5% CO_2_ for 2h. Antigens were separately diluted down to a concentration of 60nM in 50µl of the 2:1 ratio of RPMI media (Gibco) + FLIPR Calcium 6 dye (Molecular Devices) and plated in a black, clear bottom 96-well plate. The final concentration of antigen would be 30nM based on the additional 50µl of cells added during the assay. A positive control stimulant, Anti-human IgM F(ab’)_2_ (Jackson Immuno) for the IgM Ramos cell lines or Anti-human IgG F(ab’)_2_ (SouthernBiotech) for the IgG cell lines was also included in the antigen plate. Using the FlexStation 3 multi-mode microplate reader (Molecular Devices), 50µl of the cells were added to 50µl of protein or Anti-human IgM F(ab’)_2_ diluted in RPMI/dye and read continuously for 5min. The fluorescence of a blank well containing only the RPMI/dye mixture was used for background subtraction. After subtraction the antigen fluorescence was then normalized with respect to the maximum signal of the IgM or IgG control and calcium flux values were presented as a percentage. Calcium flux data are an average of at least 2 measurements.

### Cryo-EM of BCR

For cryo-EM grid preparation 2-2.5mg/ml sample concentration was used for both the BCRs. A 3.5-μL drop of protein was deposited on a Quantifoil Au-1.2/1.3 grid (Electron Microscopy Sciences, PA) that had been glow discharged for 40 seconds using a PELCO easiGlow Cleaning System (Ted Pella). After a 30 seconds incubation in >95% humidity, excess protein was blotted away for 2.5 s before being plunge frozen into liquid ethane. Data was collected on a Titan Krios 300 keV (Thermo Fisher) microscope with a K3 camera (Gatan) with a nominal defocus range of −3 to −2.8μm and 57-60 e/A^2^ electron dose range. Cryo-EM data processing was done with CryoSparc software (Punjani et al., 2017)^41^. Raw movies were motion corrected using Patch Motion Correction followed by Contrast Transfer Function (CTF) estimation. Micrographs with CTF values greater than 8 Å were discarded. Automated blob picker software was used to assign the particle position, and the particles were extracted with 432-pixel extraction box size. Following particle extraction, multiple rounds of 2D classification was performed to remove junk particles. As the BCRs were flexible, the 2D classes yielded a mix of particle views that were aligned on the Fab and Fc region (“Fab end”), the Fc, CoR and transmembrane (“TM end”) or the whole BCR. The proportions of particles showing ordered full length BCR was ∼10% of the total particle population. A total of ∼450000 particles were selected after 2D classification. After grouping these particles by the BCR region they were aligned on, ab initio 3D reconstructions were performed. Due to low particle number, the reconstruction of full length BCR did not yield optimal results. Further heterogenous refinements were performed to eliminate remaining junk particles and sort the particles into their classes, followed by the non-uniform refinements on the selected classes. The full length BCRs were reconstructed by grafting the “Fab end” and the “TM end” reconstructions by aligning them over the common Fc region. Particle subtraction followed by local refinement was used to further improve the resolution. Coordinate fitting was performed in Chimera, Coot and phenix software^42–44^. ChimeraX and Pymol was used for visualization and for making figures^45,46^.

### Molecular dynamics simulations of free BCR in POPC

We embedded the CH31 BCR structure in a phosphatidylcholine (POPC) membrane lipid bilayer and solvated the system in 0.15 M NaCl solution using the CHARMM-GUI webserver^49–52^. The resulting system size was ∼1.5 million atoms in a rectangular simulation box. The CHARMM36m force field parameter sets^53^ were used to simulate the CH31 BCR in the POPC membrane lipid bilayer. We used the default input files from the CHARMM-GUI webserver^49–52^ for GROMACS^54^ simulations of membrane proteins for our simulations. We applied periodic boundary conditions to the simulation systems and restrained bonds containing hydrogen atoms with the LINCS^55^ algorithm. We calculated the electrostatic interactions using the particle mesh Ewald (PME) summation^56^ and the Verlet cutoff scheme^57^ with a cutoff distance of 12 Å for long-range interactions. We kept the temperature constant at 310 K using the Nose-Hoover thermostat^58^ with a friction coefficient of 1.0 ps^-1^. Furthermore, we kept the pressure constant at 1.0 bar using the Parrinello-Rahman barostat^59^ with semi-isotropic coupling. We set the pressure coupling constant to 5 ps and the compressibility to 4.5×10^-5^ bar-1. The simulation systems were energetically minimized to a maximum of 5,000 steps using the steepest-descent algorithm. We applied position restraints on the backbone atoms with a force constant of 4,000 kJ.mol^-1^.nm^-2^, on the side chain atoms with a force constant of 2,000 kJ.mol^-1^.nm ^-2^, and on the lipids and dihedral angles with a force constant of 1,000 kJ.mol^-1^.nm^-2^. We then equilibrated the systems with the constant number, volume, and temperature (NVT) ensemble for a total of 375,000 steps, with a time step of 1 fs. We gradually reduced the force constants for position restraints from 4,000 to 2,000 to 1,000 kJ.mol^-1^.nm^-2^ for backbone atoms, from 2,000 to 1,000 to 500 kJ.mol^-1^.nm^-2^ for side chain atoms, from 1,000 to 400 kJ.mol^-1^.nm^-2^ for lipids, and from 1,000 to 400 to 200 kJ.mol^-1^.nm^-2^ for dihedral angles, after every 125,000 steps. We further equilibrated the systems with the constant number, pressure, and temperature (NPT) ensemble for a total of 750,000 steps, with a time step of 2 fs. We gradually reduced the force constants for position restraints from 500 to 200 to 50 kJ.mol^-1^.nm^-2^ for backbone atoms, from 200 to 50 to 0 kJ.mol^-1^.nm^-2^ for side chain atoms, from 200 to 40 to 0 kJ.mol^-1^.nm^-2^ for lipids, and from 200 to 100 to 0 kJ.mol^-1^.nm^-2^ for dihedral angles, after every 250,000 steps. Finally, we equilibrated the systems with a short 25ns conventional MD (cMD) simulation using a time step of 2 fs. We then performed five 500ns conventional MD production simulations.

After our simulations, we performed the principal component analysis (PCA) to obtain the important low-energy conformational states observed for the CH31 BCR (**Fig. S12A-S12C**). We carried out the PCA following the protocols described in the AmberTools^60,61^ tutorial. Furthermore, we used the CPPTRAJ^62^ simulation analysis tool and the GROMACS 2022^54^ simulation package to calculate the root-mean-square fluctuation (RMSF) from each simulation replica and averaged them to find the average RMSF for the CH31 BCR (**Fig. S12D-S12E**).

## Supplementary Materials

Fig. S1. Ramos cell line phenotypic data and activation with control stimulant, [Relates to Fig.1 and Methods].

Fig. S2. Representative NSEM analysis of purified BCR from Ramos cells [Relates to Figure # 1 and Methods].

Fig. S3. SPR Curves and Fits for 2G12.

Fig. S4. SPR Curves and Fits for CH31.

Fig. S5. Activation of Ramos Cells expressing IgM or IgG BCRs with specificity of CH31 to Env trimers and trimer multimers.

Fig. S6. Cryo-EM data processing for 2G12 IgG BCR.

Fig. S7. Local resolution estimation and local refinement 2G12 IgG BCR

Fig. S8. Igα drives class-specific configuration of IgG versus IgM BCR.

Fig. S9. Comparison of 2G12 IgG BCR with 7WSO

Fig. S10. Cryo-EM data processing for CH31 IgM BCR

Fig. S11. Interaction of co-receptors with Fc domains

Fig. S12. Dynamics of the BCR in POPC lipid bilayer obtained from the MD simulations.

Table S1. Kinetic Rates Measurements for 2G12 [Relates to Main Figure 2].

Table S2. Kinetic Rates Measurements for CH31.

Table S3. Comparison of the maximum calcium flux (Ca2+ Flux) responses and activation of Ramos cells expressing 2G12 IgM or IgG BCRS with Env trimers and multimers [Relates to Main Figure 3]

Table S4. 2G12 IgG and CH31 IgM BCR Cryo-EM data collection and refinement statistics.

Movie S1. Hinge flexibility defines dynamic positioning of 2G12 Fab in the BCR.

Movie S2. Dynamic of the CH31 BCR observed for atomistic molecular dynamics simulations.

## Acknowledgments

We thank Dr. Bart Haynes, DHVI, Director of CHAVD (Duke Consortia) for providing facility resources, scientific advice, and critical comments. We thank Kevin Saunders, Elizabeth Donahue (Duke HVI) for expressing and purifying Env gp120 and gp140 trimers and NPs. We thank the Duke CHSB U54 program management team members (Aaron Cook, Program Manager; Heather Brown, Grant Manager; Angela Burnette, Regulatory Coordinator; Yunfei Wang, Biostatistician; Amber (Kelly) Wright, Communications), DHVI teams from BIAcore Facility, Flow cytometry, Protein Expression (Kevin Saunders) Center and the DHVI Finance and administrative teams for their support. We are grateful to Rogier W. Sanders, Ronald Derking, and Tom Bijl (Amsterdam UMC) for the expression and production of GT1.2 gp140 trimer and GT1.2 NPs.

## Author contributions

Conceptualization: SMA. Methodology: BT, JA, KC, KA, AH, APK, PP, RJE, KS, SG, PA, SMA. Investigation: BT, JA, KC, PA, KA, APK, PP, KM, TS, HD, RJE, KJ, ML. Supervision: SMA, PA, KC, RJE. Writing – original draft: SMA, PA, BT, JA. Writing – review & editing: all authors. Funding acquisition: SMA, PA, BFH. Project administration: AC.

## Data and materials availability

All data analysis needed to evaluate the conclusions in the paper are present in the paper or the Supplementary Materials.

## REFERENCES

1 McHeyzer-Williams, L. J. & McHeyzer-Williams, M. G. Antigen-specific memory B cell development. Annu Rev Immunol 23, 487–513 (2005). 10.1146/annurev.immunol.23.021704.115732

2 Schamel, W. W. & Reth, M. Monomeric and oligomeric complexes of the B cell antigen receptor. Immunity 13, 5–14 (2000). 10.1016/s1074-7613(00)00003-0

3 Reth, M. Antigen receptors on B lymphocytes. Annu Rev Immunol 10, 97–121 (1992). 10.1146/annurev.iy.10.040192.000525

4 Reth, M. Antigen receptor tail clue. Nature 338, 383–384 (1989). 10.1038/338383b0

5 Flaswinkel, H. & Reth, M. Dual role of the tyrosine activation motif of the Ig-alpha protein during signal transduction via the B cell antigen receptor. Embo j 13, 83–89 (1994). 10.1002/j.1460-2075.1994.tb06237.x

6 Tolar, P., Sohn, H. W., Liu, W. & Pierce, S. K. The molecular assembly and organization of signaling active B-cell receptor oligomers. Immunol Rev 232, 34–41 (2009). 10.1111/j.1600-065X.2009.00833.x

7 Pierce, S. K. & Liu, W. The tipping points in the initiation of B cell signalling: how small changes make big differences. Nat Rev Immunol 10, 767–777 (2010). 10.1038/nri2853

8 Tolar, P. & Pierce, S. K. A conformation-induced oligomerization model for B cell receptor microclustering and signaling. Curr Top Microbiol Immunol 340, 155–169 (2010). 10.1007/978-3-642-03858-7_8

9 Ma, X. et al. Cryo-EM structures of two human B cell receptor isotypes. Science 377, 880–885 (2022). 10.1126/science.abo3828

10 Su, Q. et al. Cryo-EM structure of the human IgM B cell receptor. Science 377, 875–880 (2022). 10.1126/science.abo3923

11 Dong, Y. et al. Structural principles of B cell antigen receptor assembly. Nature 612, 156–161 (2022). 10.1038/s41586-022-05412-7

12 Tolar, P., Sohn, H. W. & Pierce, S. K. The initiation of antigen-induced B cell antigen receptor signaling viewed in living cells by fluorescence resonance energy transfer. Nat Immunol 6, 1168–1176 (2005). 10.1038/ni1262

13 Calarese, D. A. et al. Antibody domain exchange is an immunological solution to carbohydrate cluster recognition. Science 300, 2065–2071 (2003). 10.1126/science.1083182

14 Doores, K. J. et al. Envelope glycans of immunodeficiency virions are almost entirely oligomannose antigens. Proc Natl Acad Sci U S A 107, 13800–13805 (2010). 10.1073/pnas.1006498107

15 Acharya, P. et al. A glycan cluster on the SARS-CoV-2 spike ectodomain is recognized by Fab-dimerized glycan-reactive antibodies. bioRxiv (2020). 10.1101/2020.06.30.178897

16 Scanlan, C. N. et al. The broadly neutralizing anti-human immunodeficiency virus type 1 antibody 2G12 recognizes a cluster of alpha1-->2 mannose residues on the outer face of gp120. J Virol 76, 7306–7321 (2002). 10.1128/jvi.76.14.7306-7321.2002

17 Williams, W. B. et al. Fab-dimerized glycan-reactive antibodies are a structural category of natural antibodies. Cell 184, 2955–2972.e2925 (2021). 10.1016/j.cell.2021.04.042

18 Scanlan, C. N. et al. The carbohydrate epitope of the neutralizing anti-HIV-1 antibody 2G12. Adv Exp Med Biol 535, 205–218 (2003). 10.1007/978-1-4615-0065-0_13

19 Sanders, R. W. et al. The mannose-dependent epitope for neutralizing antibody 2G12 on human immunodeficiency virus type 1 glycoprotein gp120. J Virol 76, 7293–7305 (2002). 10.1128/jvi.76.14.7293-7305.2002

20 Trkola, A. et al. Human monoclonal antibody 2G12 defines a distinctive neutralization epitope on the gp120 glycoprotein of human immunodeficiency virus type 1. J Virol 70, 1100–1108 (1996). 10.1128/jvi.70.2.1100-1108.1996

21 Murin, C. D. et al. Structure of 2G12 Fab2 in complex with soluble and fully glycosylated HIV-1 Env by negative-stain single-particle electron microscopy. J Virol 88, 10177–10188 (2014). 10.1128/jvi.01229-14

22 Doores, K. J., Fulton, Z., Huber, M., Wilson, I. A. & Burton, D. R. Antibody 2G12 recognizes di-mannose equivalently in domain- and nondomain-exchanged forms but only binds the HIV-1 glycan shield if domain exchanged. J Virol 84, 10690–10699 (2010). 10.1128/JVI.01110-10

23 Doores, K. J. et al. 2G12-expressing B cell lines may aid in HIV carbohydrate vaccine design strategies. J Virol 87, 2234–2241 (2013). 10.1128/JVI.02820-12

24 Bonsignori, M. et al. Two distinct broadly neutralizing antibody specificities of different clonal lineages in a single HIV-1-infected donor: implications for vaccine design. J Virol 86, 4688–4692 (2012). 10.1128/jvi.07163-11

25 Bonsignori, M. et al. Inference of the HIV-1 VRC01 Antibody Lineage Unmutated Common Ancestor Reveals Alternative Pathways to Overcome a Key Glycan Barrier. Immunity 49, 1162–1174.e1168 (2018). 10.1016/j.immuni.2018.10.015

26 Zhou, T. et al. Structural basis for broad and potent neutralization of HIV-1 by antibody VRC01. Science 329, 811–817 (2010). 10.1126/science.1192819

27 Zhou, T. et al. Multidonor analysis reveals structural elements, genetic determinants, and maturation pathway for HIV-1 neutralization by VRC01-class antibodies. Immunity 39, 245–258 (2013). 10.1016/j.immuni.2013.04.012

28 Hossain, M. A. et al. B cells expressing IgM B cell receptors of HIV-1 neutralizing antibodies discriminate antigen affinities by sensing binding association rates. Cell Rep 39, 111021 (2022). 10.1016/j.celrep.2022.111021

29 Ortiz, Y. et al. The CH1 domain influences the expression and antigen sensing of the HIV-specific CH31 IgM-BCR and IgG-BCR. Proc Natl Acad Sci U S A 121, e2404728121 (2024). 10.1073/pnas.2404728121

30 Henderson, R. et al. Selection of immunoglobulin elbow region mutations impacts interdomain conformational flexibility in HIV-1 broadly neutralizing antibodies. Nat Commun 10, 654 (2019). 10.1038/s41467-019-08415-7

31 Stanfield, R. L., Zemla, A., Wilson, I. A. & Rupp, B. Antibody elbow angles are influenced by their light chain class. J Mol Biol 357, 1566–1574 (2006). 10.1016/j.jmb.2006.01.023

32 Lesk, A. M. & Chothia, C. Elbow motion in the immunoglobulins involves a molecular ball-and-socket joint. Nature 335, 188–190 (1988). 10.1038/335188a0

33 Rothlisberger, D., Honegger, A. & Pluckthun, A. Domain interactions in the Fab fragment: a comparative evaluation of the single-chain Fv and Fab format engineered with variable domains of different stability. J Mol Biol 347, 773–789 (2005). 10.1016/j.jmb.2005.01.053

34 Do, H. N., Zhao, M., Alam, S.M. Gnanakaran, S. Dynamics and activation of membrane-bound B cell receptor assembly bioRxiv (2024). doi: 10.1101/2024.07.10.602784;

35 Yang, J. & Reth, M. Oligomeric organization of the B-cell antigen receptor on resting cells. Nature 467, 465–469 (2010). 10.1038/nature09357

36 Reth, M. Discovering immunoreceptor coupling and organization motifs. Front Immunol 14, 1253412 (2023). 10.3389/fimmu.2023.1253412

37 Veneziano, R. et al. Role of nanoscale antigen organization on B-cell activation probed using DNA origami. Nat Nanotechnol 15, 716–723 (2020). 10.1038/s41565-020-0719-0

38 Maity, P. C. et al. B cell antigen receptors of the IgM and IgD classes are clustered in different protein islands that are altered during B cell activation. Sci Signal 8, ra93 (2015). 10.1126/scisignal.2005887

39 Benjamin, D. et al. Immunoglobulin secretion by cell lines derived from African and American undifferentiated lymphomas of Burkitt’s and non-Burkitt’s type. J Immunol 129, 1336–1342 (1982).

40 Zivanov, J. et al. New tools for automated high-resolution cryo-EM structure determination in RELION-3. Elife 7 (2018). 10.7554/eLife.42166

41 Punjani, A., Rubinstein, J. L., Fleet, D. J. & Brubaker, M. A. cryoSPARC: algorithms for rapid unsupervised cryo-EM structure determination. Nature methods 14, 290–296 (2017).

42 Pettersen, E. F. et al. UCSF Chimera—a visualization system for exploratory research and analysis. Journal of computational chemistry 25, 1605–1612 (2004).

43 Emsley, P., Lohkamp, B., Scott, W. G. & Cowtan, K. Features and development of Coot. Acta Crystallographica Section D: Biological Crystallography 66, 486–501 (2010).

44 Afonine, P. V. et al. Real-space refinement in PHENIX for cryo-EM and crystallography. Acta Crystallographica Section D: Structural Biology 74, 531–544 (2018).

45 Goddard, T. D. et al. UCSF ChimeraX: Meeting modern challenges in visualization and analysis. Protein science 27, 14–25 (2018).

46 Schrödinger, L. The PyMOL Molecular Graphics System, Version 1.8. (No Title) (2015).

47 Bianchi, M et al; Electron-Microscopy-Based Epitope Mapping Defines Specificities of Polyclonal Antibodies Elicited during HIV-1 BG505 Envelope Trimer Immunization,Immunity,Volume 49, Issue 2,2018,Pages 288-300.e8,ISSN 1074-7613,10.1016/j.immuni.2018.07.009.

48 Luo, X.M., Maarschalk, E., O’Connell, R.M., Wang, P., Yang, L., and Baltimore, D. (2009). Engineering human hematopoietic stem/progenitor cells to produce a broadly neutralizing anti-HIV antibody after in vitro maturation to human B lymphocytes. Blood 113, 1422–1431.

49 S. Jo, T. Kim, V. Iyer, W. Im, CHARMM-GUI: A Web-based Graphical User Interface for CHARMM. Journal of Computational Chemistry 29, 1859–1865 (2008).

50 E. Wu et al., CHARMM-GUI Membrane Builder Toward Realistic Biological Membrane Simulations. Journal of Computational Chemistry 35, 1997–2004 (2014).

51 S. Jo, J. Lim, J. Klauda, W. Im, CHARMM-GUI Membrane Builder for Mixed Bilayers and Its Application to Yeast Membranes. Biophysical Journal 97, 50–58 (2009).

52 J. Lee et al., CHARMM-GUI Input Generator for NAMD, GROMACS, AMBER, OpenMM, and CHARMM/OpenMM Simulations using the CHARMM36 Additive Force Field. Journal of Chemical Theory and Computation 12, 405-413 (2016).

53 J. Huang et al., CHARMM36m: an improved force field for folded and intrinscially disordered proteins. Nature Methods 14, 71–73 (2017).

54 M. J. Abraham et al., GROMACS: High performance molecular simulations through multi-level parallelism from laptops to supercomputers. SoftwareX 1–2, 19-25 (2015).

55 B. Hess, H. Bekker, H. J. C. Berendsen, J. G. E. M. Fraaije, LINCS: A linear constraint solver for molecular simulations. Journal of Computational Chemistry 18, 1463–1472 (1998).

56 U. Essmann et al., A Smooth Particle Mesh Ewald Method. Journal of Chemical Physics 103, (1995).

57 H. Grubmuller, H. Heller, A. Windemuth, K. Schulten, Generalized Verlet Algorithm for Efficient Molecular Dynamics Simulations with Long-Range Interactions. Molecular Simulation 6, 121–142 (1991).

58 D. J. Evans, B. L. Holian, The Nose-Hoover thermostat. The Journal of Chemical Physics 83, 4069–4074 (1985).

59 M. Parrinello, A. Rahman, Polymorphic Transitions in Single Crystals: A New Molecular Dynamics Method. Journal of Applied Physics 52, 7182–7190 (1981).

60 D. A. Case et al., AmberTools. Journal of Chemical Information and Modeling 63, 6183–6191 (2023).

61. D. A. Case et al., AMBER 2020. (2020).

62 D. R. Roe, I. T. E. Cheatham, PTRAJ and CPPTRAJ: software for processing and analysis of molecular dynamics trajectory data. Journal of Chemical Theory and Computation 9, 3084–3095 (2013).

