## Supplemental Figures for "Anti-HIV-1 B cell antigen receptor signaling and structure"

### Supplementary Material


1. **Fig. S1. Ramos cell line phenotypic data and activation with control stimulant [Relates to Fig.1 and Methods]**
2. **Fig. S2. Representative NSEM analysis of purified BCR from Ramos cells [Relates to Figure # 1 and Methods].**
3. **Fig. S3. SPR Curves and Fits for 2G12.**
4. **Table S1. Kinetic Rates Measurements for 2G12 [Relates to Main Figure 2].**
5. **Fig. S4. SPR Curves and Fits for CH31.**
6. **Table S2. Kinetic Rates Measurements for CH31**
7. **Fig. S5. Activation of Ramos Cells expressing IgM or IgG BCRs with specificity of CH31 to Env trimers and trimer multimers.**
8. **Table S3. Comparison of the maximum calcium flux (Ca2+ Flux) responses and activation of Ramos cells expressing 2G12 IgM or IgG BCRS with Env trimers and multimers [Relates to Main Figure 3]**
9. **Fig. S6. Cryo-EM data processing for 2G12 IgG BCR.**
10. **Fig. S7. Local resolution estimation and local refinement 2G12 IgG BCR**
11. **Fig. S8. Igα drives class-specific configuration of IgG versus IgM BCR.**
12. **Fig. S9. Comparison of 2G12 IgG BCR with 7WSO**
13. **Fig. S10. Cryo-EM data processing for CH31 IgM BCR**
14. **Fig. S11. Interaction of co-receptors with Fc domains**
15. **Table S4. 2G12 IgG and CH31 IgM BCR Cryo-EM data collection and refinement statistics.**
16. **Fig. S12. Dynamics of the BCR in POPC lipid bilayer obtained from the MD simulations**
17. **Movie S1. Hinge flexibility defines dynamic positioning of 2G12 Fab in the BCR.**
18. **Movie S2. Dynamic of the CH31 BCR observed for atomistic molecular dynamics simulations**

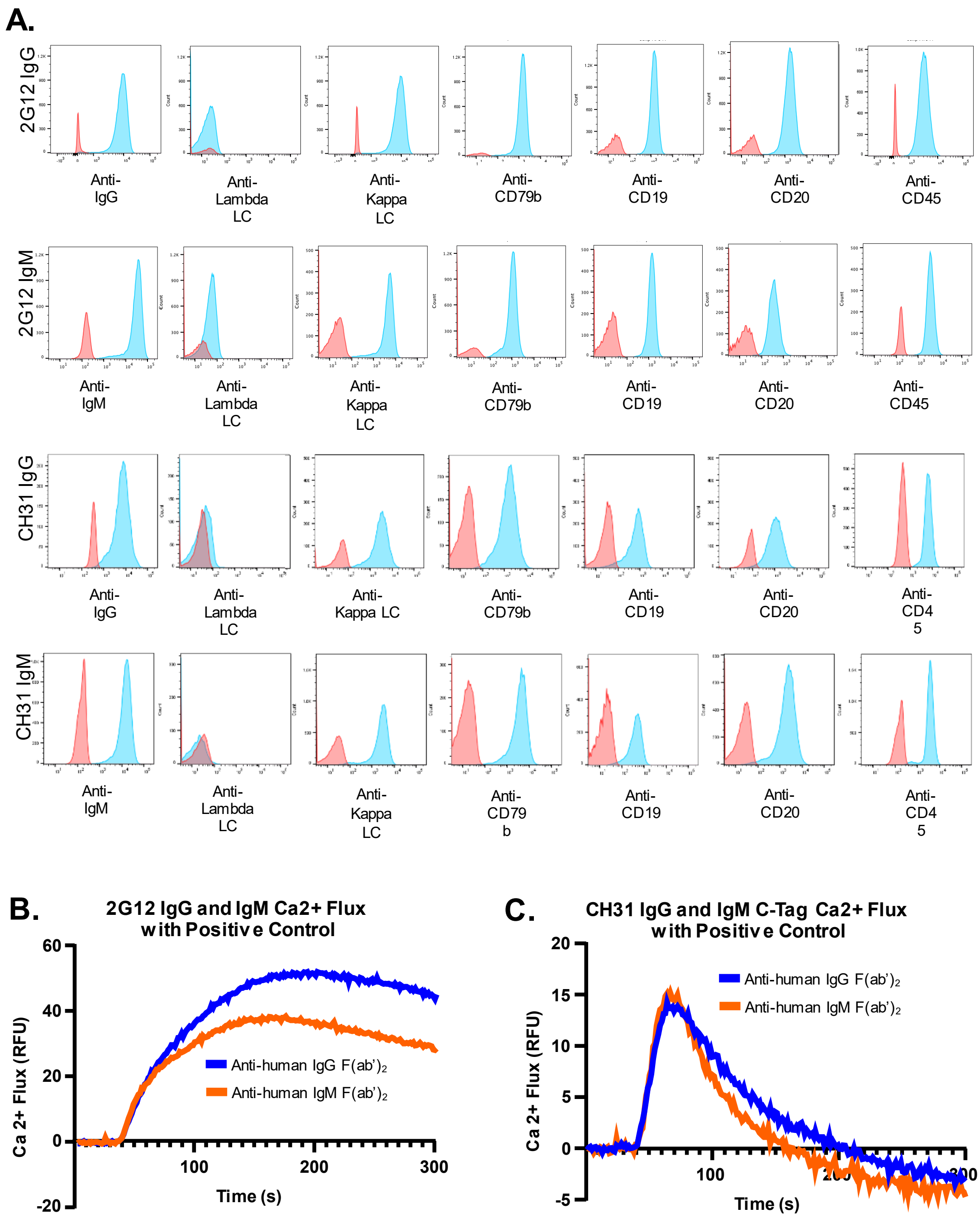

A.
2G12 IgG
Anti-IgG
Anti-Lambda LC
Anti-Kappa LC
Anti-CD79b
Anti-CD19
Anti-CD20
Anti-CD45
2G12 IgM
Anti-IgM
Anti-Lambda LC
Anti-Kappa LC
Anti-CD79b
Anti-CD19
Anti-CD20
Anti-CD45
CH31 IgG
Anti-IgG
Anti-Lambda LC
Anti-Kappa LC
Anti-CD79b
Anti-CD19
Anti-CD20
Anti-CD45
CH31 IgM
Anti-IgM
Anti-Lambda LC
Anti-Kappa LC
Anti-CD79b
Anti-CD19
Anti-CD20
Anti-CD45
B.
2G12 IgG and IgM Ca2+ Flux with Positive Control
Ca 2+ Flux (RFU)
Time (s)
60
40
20
0
-20
100
200
300
Anti-human IgG F(ab')2
Anti-human IgM F(ab')2
C.
CH31 IgG and IgM C-Tag Ca2+ Flux with Positive Control
Ca 2+ Flux (RFU)
Time (s)
20
15
10
5
0
-5
100
Anti-human IgG F(ab')2
Anti-human IgM F(ab')2

**Fig. S1. Ramos cell line phenotypic data and activation with control stimulant [Relates to Fig.1 and Methods]. (A)** Ramos cell lines expressing 2G12 IgM, 2G12 IgG, CH31 IgM AID KO, and CH31 IgG AID KO BCR were testing using flow cytometry for BCR and surface markers expression levels. All cell lines (blue) showed high expression for IgM/IgG and kappa light chain when compared to unstained cells (red). All cell lines showed expression of CD79b, CD19, CD20, and CD45. **(B)** Calcium flux of Ramos cells expressing BCRs of 2G12 IgG (blue) and IgM (orange) by the positive control Anti-human F(ab')2 at concentrations of 25ug/mL for the anti-IgG antibody and at 50ug/mL for the anti-IgM antibody. **(C)** Calcium flux of Ramos cells expressing BCRs of CH31 IgG (blue) and IgM (orange) by the positive control Anti-human F(ab')2 at concentrations of 25ug/mL for the anti-IgG antibody and at 50ug/mL for the anti-IgM antibody. Data shown here for the CH31 Ramos cell line with AID-/-. All Ramos cell lines were treated with the anti-human F(ab')2 along with the FLIPR Calcium 6 dye and monitored for 5 min using the Flexstation3 plate reader. The fluorescence of a blank well containing only RPMI/dye was used for background subtraction. The calcium flux profiles are an averaged of at least three measurements and demonstrate the functionality of the three cell lines.

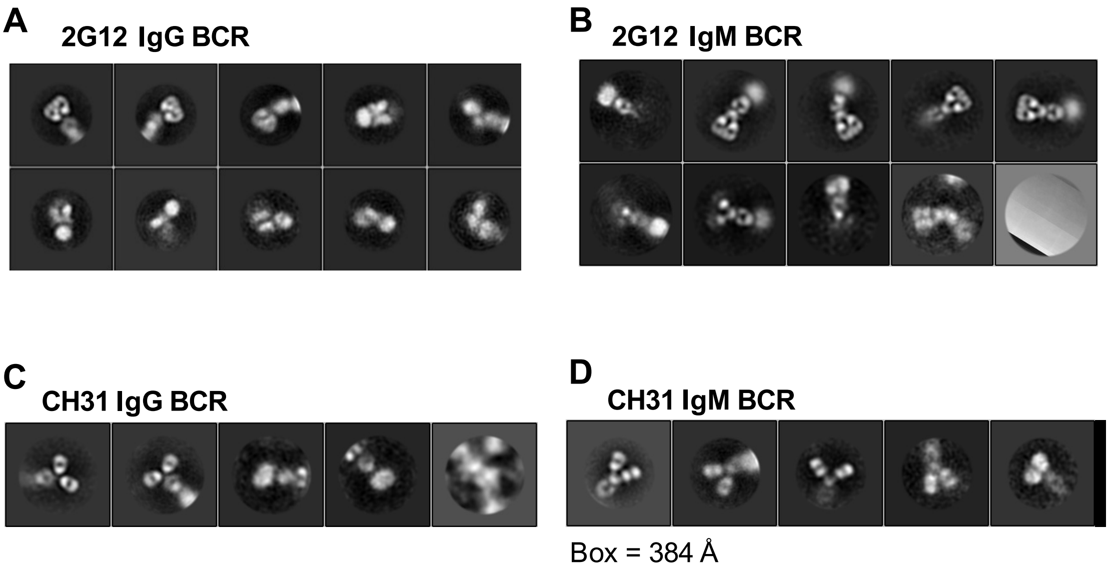


**Fig. S2. Representative NSEM analysis of purified BCR from Ramos cells [Relates to Figure # 1 and Methods].** **(A)** NSEM 2D class averages of 2G12 IgG BCR showing predominantly I-shaped configuration; particles picked in final round = 12,000 **(B)** NSEM 2D class averages of 2G12 IgM BCR showing predominantly I-shaped configuration; particles picked in final round = 16,040. **(C)** NSEM 2D class averages of CH31 IgG BCR showing predominantly Y-shaped configuration; particles picked in final round = 4,955. **(D)** NSEM 2D class averages of CH31 IgM BCR showing predominantly Y-shaped configuration; particles picked in final round = 8,922. Grids were examined on a Philips EM420 electron microscope operating at 120 kV and nominal magnification of 49,000x, and images were collected on a 76 Mpix CCD camera at 2.4 Å/pixel. Images were analyzed by 2D class averages using standard protocols with Relion 3.0.

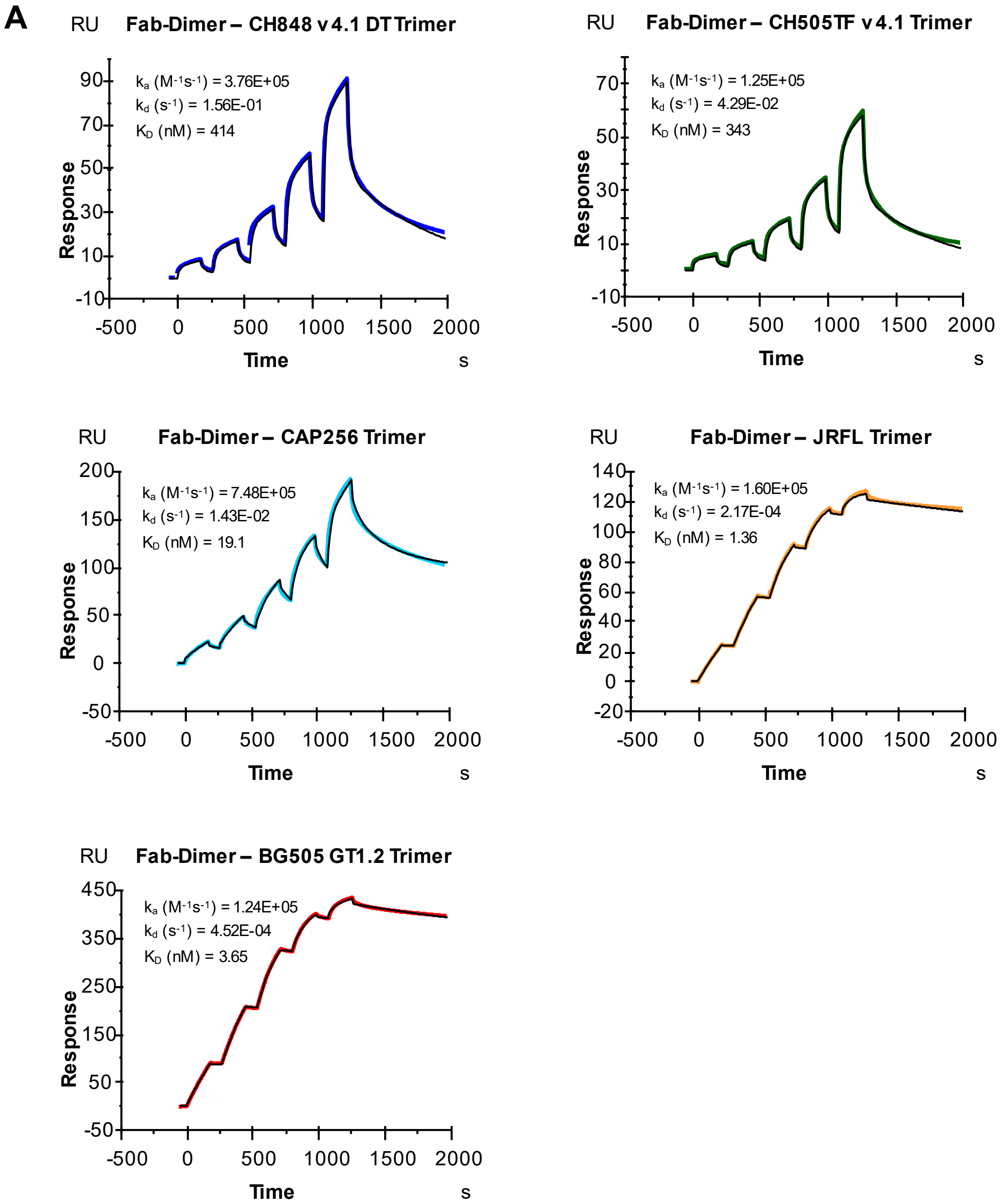
A
Fab-Dimer – CH848 v4.1 DT Trimer
ka (M-1s-1) = 3.76E+05
kd (s-1) = 1.56E-01
KD (nM) = 414
Fab-Dimer – CH505TF v4.1 Trimer
ka (M-1s-1) = 1.25E+05
kd (s-1) = 4.29E-02
KD (nM) = 343
Fab-Dimer – CAP256 Trimer
ka (M-1s-1) = 7.48E+05
kd (s-1) = 1.43E-02
KD (nM) = 19.1
Fab-Dimer – JRFL Trimer
ka (M-1s-1) = 1.60E+05
kd (s-1) = 2.17E-04
KD (nM) = 1.36
Fab-Dimer – BG505 GT1.2 Trimer
ka (M-1s-1) = 1.24E+05
kd (s-1) = 4.52E-04
KD (nM) = 3.65
RU
Response
Time
s

**B**

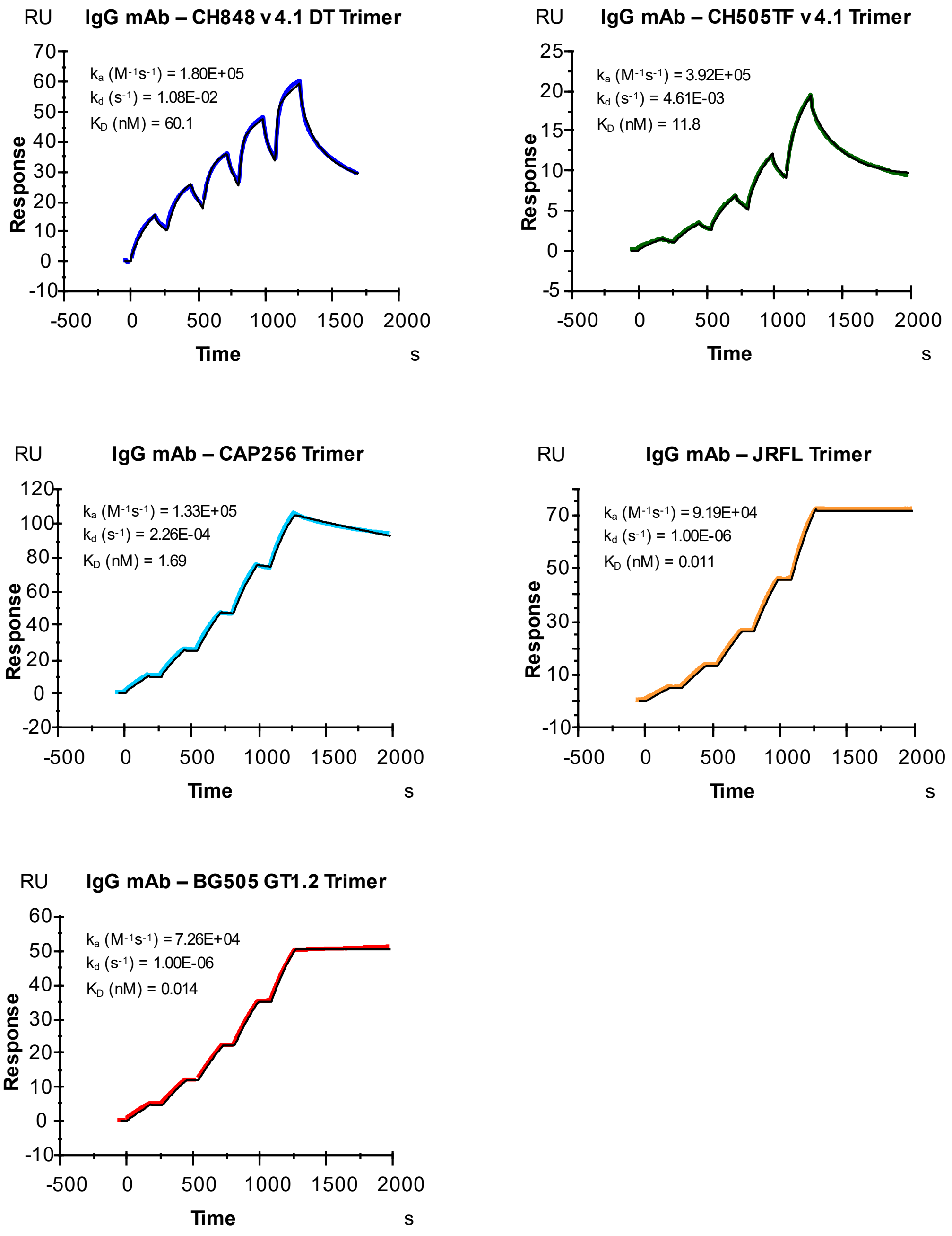
IgG mAb – CH848 v4.1 DT Trimer
RU
ka (M-1s-1) = 1.80E+05
kd (s-1) = 1.08E-02
KD (nM) = 60.1
Response
Time
s
IgG mAb – CH505TF v4.1 Trimer
RU
ka (M-1s-1) = 3.92E+05
kd (s-1) = 4.61E-03
KD (nM) = 11.8
Response
Time
s
IgG mAb – CAP256 Trimer
RU
ka (M-1s-1) = 1.33E+05
kd (s-1) = 2.26E-04
KD (nM) = 1.69
Response
Time
s
IgG mAb – JRFL Trimer
RU
ka (M-1s-1) = 9.19E+04
kd (s-1) = 1.00E-06
KD (nM) = 0.011
Response
Time
s
IgG mAb – BG505 GT1.2 Trimer
RU
ka (M-1s-1) = 7.26E+04
kd (s-1) = 1.00E-06
KD (nM) = 0.014
Response
Time
s

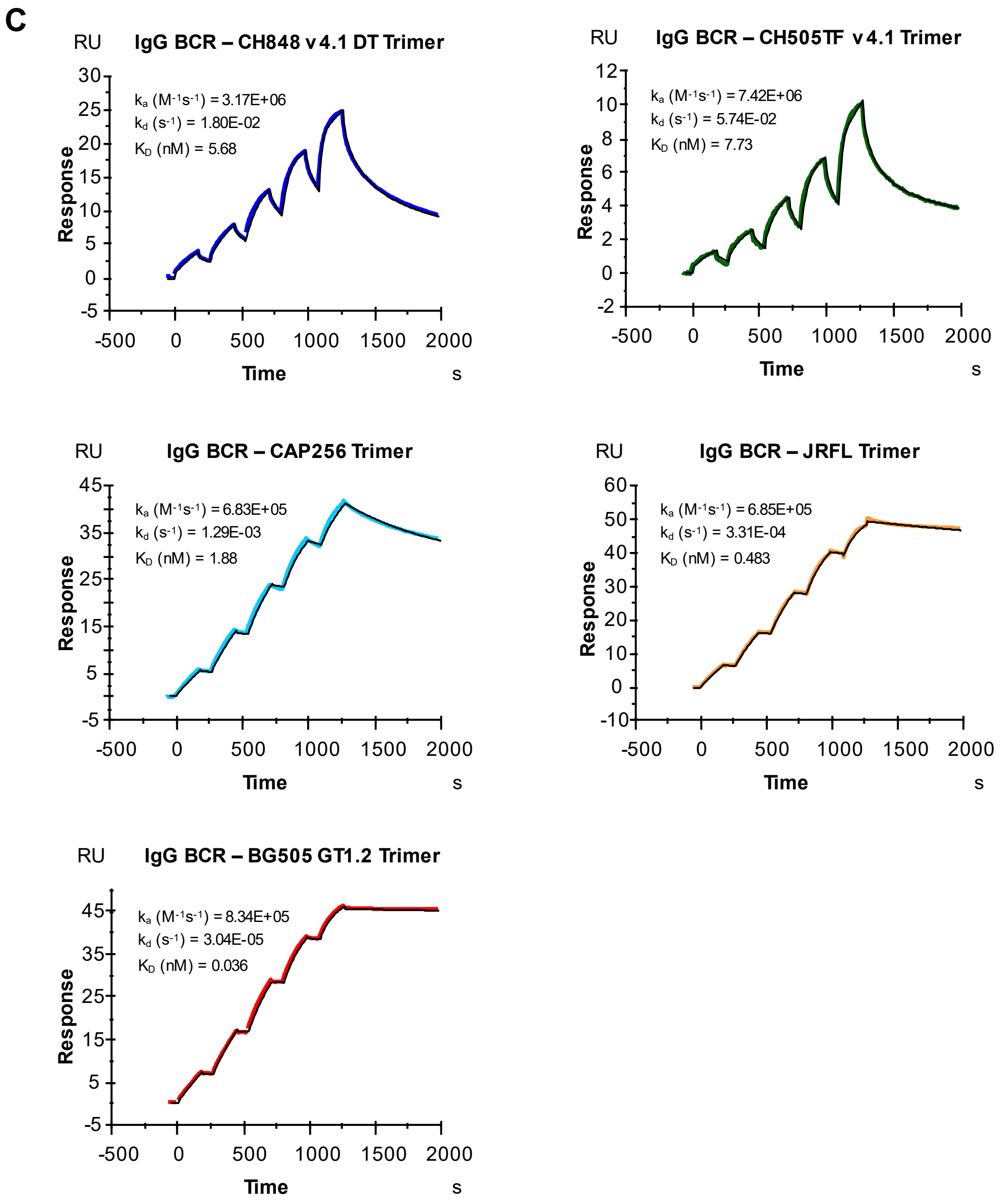
C
IgG BCR – CH848 v4.1 DT Trimer
k_a (M^-1s^-1) = 3.17E+06
k_d (s^-1) = 1.80E-02
K_D (nM) = 5.68
IgG BCR – CH505TF v4.1 Trimer
k_a (M^-1s^-1) = 7.42E+06
k_d (s^-1) = 5.74E-02
K_D (nM) = 7.73
IgG BCR – CAP256 Trimer
k_a (M^-1s^-1) = 6.83E+05
k_d (s^-1) = 1.29E-03
K_D (nM) = 1.88
IgG BCR – JRFL Trimer
k_a (M^-1s^-1) = 6.85E+05
k_d (s^-1) = 3.31E-04
K_D (nM) = 0.483
IgG BCR – BG505 GT1.2 Trimer
k_a (M^-1s^-1) = 8.34E+05
k_d (s^-1) = 3.04E-05
K_D (nM) = 0.036
RU
Response
Time
s

D

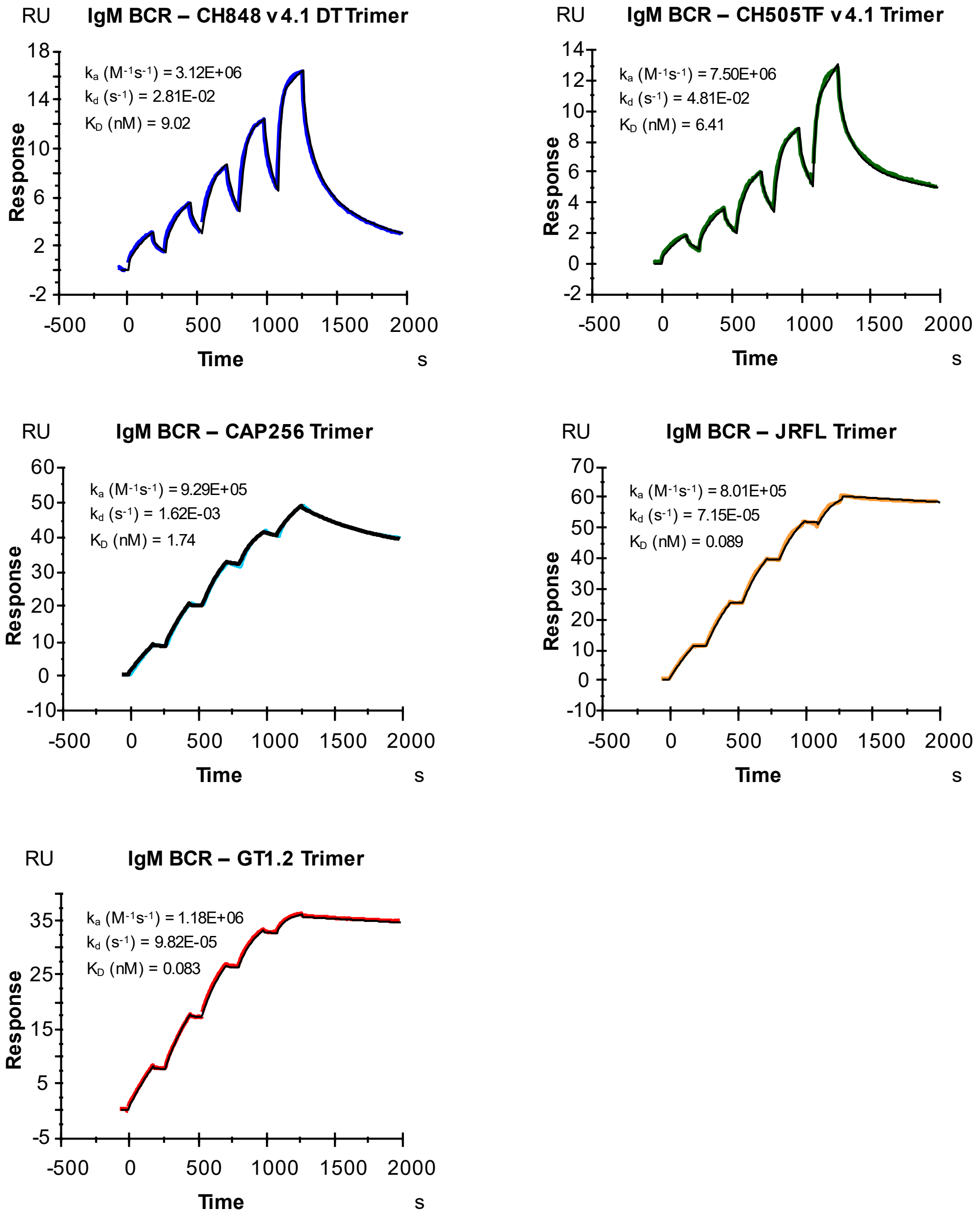
IgM BCR – CH848 v4.1 DT Trimer
RU
Response
Time
s
$k_a$ ($M^{-1}s^{-1}$) = 3.12E+06
$k_d$ ($s^{-1}$) = 2.81E-02
$K_D$ (nM) = 9.02
IgM BCR – CH505TF v4.1 Trimer
RU
Response
Time
s
$k_a$ ($M^{-1}s^{-1}$) = 7.50E+06
$k_d$ ($s^{-1}$) = 4.81E-02
$K_D$ (nM) = 6.41
IgM BCR – CAP256 Trimer
RU
Response
Time
s
$k_a$ ($M^{-1}s^{-1}$) = 9.29E+05
$k_d$ ($s^{-1}$) = 1.62E-03
$K_D$ (nM) = 1.74
IgM BCR – JRFL Trimer
RU
Response
Time
s
$k_a$ ($M^{-1}s^{-1}$) = 8.01E+05
$k_d$ ($s^{-1}$) = 7.15E-05
$K_D$ (nM) = 0.089
IgM BCR – GT1.2 Trimer
RU
Response
Time
s
$k_a$ ($M^{-1}s^{-1}$) = 1.18E+06
$k_d$ ($s^{-1}$) = 9.82E-05
$K_D$ (nM) = 0.083

**Fig. S3. SPR Curves and Fits for 2G12. (A)** 2G12 Fab-Dimer flowed over biotinylated CH848 v4.1 DT, CH505TF v4.1, CAP256, and JRFL Env trimers immobilized via streptavidin, and His-tagged BG505 GT1.2 Env trimer captured to chip surface. **(B)** Non-tagged (without biotin or His) Env trimers as in **A** flowed over Anti-Human Fc captured 2G12 IgG mAb. **(C)** Non-tagged Env trimers as in **A** flowed over captured purified 2G12 IgG BCR. **(D)** Non-tagged Env trimers as in **A** flowed over captured purified 2G12 IgM BCR. Surface Plasmon Resonance (SPR) was used to conduct titrations with Env trimers as described in methods section. 2G12 Fab-Dimer was diluted to 12.5 – 400 nM. The following non-biotinylated Env trimers were used at the listed concentration ranges: CH505TF (2 – 200 nM), CH848 (10 – 1000 nM), GT1.2 (2 – 48 nM), CAP256 (1.5 – 48 nM), JRFL (2 – 48 nM). Heterogenous ligand model was used for biphasic curves. Plots shown are representative of at least three independent measurements with the exception of Fab-Dimer to BG505 GT1.2 being a single experiment.

| INTERACTION | | DATA FROM CURVE FITTING | | | | | MEAN | | | STANDARD ERROR | | |
|---|---|---|---|---|---|---|---|---|---|---|---|---|
| **Ligand:** | **Analyte:** | **ka1 (1/Ms)** | **ka2 (1/Ms)** | **kd1 (1/s)** | **kd2 (1/s)** | **KD (nM)** | **ka (1/Ms)** | **kd (1/s)** | **KD (nM)** | **ka (1/Ms)** | **kd (1/s)** | **KD (nM)** |
| Biotinylated CH848 v4.1 DT | 2G12 Fab-Dimer | **3.76E+05** | 9.57E+03 | **1.56E-01** | 1.39E-03 | 414 | 2.49E+05 | 1.01E-01 | 404 | 6.4E+04 | 2.8E-02 | 152 |
| | | **1.86E+05** | 6.80E+03 | **6.94E-02** | 1.32E-03 | 374 | | | | | | |
| | | **1.85E+05** | 6.90E+03 | **7.64E-02** | 1.32E-03 | 414 | | | | | | |
| Biotinylated CH505TF v4.1 | | **1.25E+05** | 1.16E+04 | **4.85E-02** | 1.77E-03 | 388 | 1.11E+05 | 4.94E-02 | 445 | 6.5E+03 | 4.7E-03 | 49 |
| | | **1.25E+05** | 1.36E+04 | **4.29E-02** | 1.65E-03 | 343 | | | | | | |
| | | **1.07E+05** | 1.28E+04 | **4.34E-02** | 1.81E-03 | 407 | | | | | | |
| | | **9.07E+04** | 1.79E+04 | **6.77E-02** | 2.23E-03 | 746 | | | | | | |
| | | **1.08E+05** | 1.57E+04 | **4.45E-02** | 1.63E-03 | 412 | | | | | | |
| Biotinylated CAP256 | | **3.36E+05** | 1.14E+03 | **9.27E-03** | 2.66E-04 | 27.6 | 5.87E+05 | 9.07E-03 | 15.4 | 9.2E+04 | 1.4E-03 | 3.4 |
| | | **7.48E+05** | 1.50E+04 | **1.43E-02** | <1.00E-06 | 19.1 | | | | | | |
| | | **5.42E+05** | 2.26E+04 | **5.18E-03** | <1.00E-06 | 9.55 | | | | | | |
| | | **9.44E+05** | 3.08E+04 | **1.15E-02** | 2.38E-04 | 12.2 | | | | | | |
| | | **3.95E+05** | 2.28E+04 | **6.03E-03** | <1.00E-06 | 15.3 | | | | | | |
| | | **5.56E+05** | 2.33E+04 | **8.12E-03** | <1.00E-06 | 14.6 | | | | | | |
| Biotinylated JRFL | | **5.10E+04** | 2.43E+04 | **1.08E-04** | <1.00E-06 | 2.11 | 1.60E+05 | 1.45E-04 | 0.901 | 2.7E+04 | 1.6E-05 | 0.185 |
| | | **1.60E+05** | 5.93E+04 | **2.17E-04** | <1.00E-06 | 1.36 | | | | | | |
| | | **1.84E+05** | 7.26E+04 | **1.52E-04** | <1.00E-06 | 0.825 | | | | | | |
| | | **1.56E+05** | 6.64E+04 | **1.14E-04** | <1.00E-06 | 0.727 | | | | | | |
| | | **1.51E+05** | 6.20E+04 | <1.00E-06 | **1.49E-04** | 0.987 | | | | | | |
| | | **2.60E+05** | 7.26E+04 | <1.00E-06 | **1.28E-04** | 0.491 | | | | | | |
| His-tagged BG505 GT1.2 | | **1.24E+05** | 4.27E+04 | <1.00E-06 | **4.52E-04** | 3.65 | 1.24E+05 | 4.52E-04 | 3.65 | N/A | N/A | N/A |
| 2G12 IgG mAb | CH848 v4.1 DT | **1.80E+05** | 4.33E+03 | **1.08E-02** | <1.00E-06 | 60.1 | 1.60E+05 | 7.85E-03 | 49.1 | 1.6E+04 | 2.9E-03 | 18.8 |
| | | **1.71E+05** | 4.05E+03 | **1.07E-02** | 2.26E-06 | 62.7 | | | | | | |
| | | **1.29E+05** | 1.39E+04 | **2.03E-03** | <1.00E-06 | 15.7 | | | | | | |
| | CH505TF v4.1 | **3.92E+05** | 3.56E+04 | **4.61E-03** | <1.00E-06 | 11.8 | 4.33E+05 | 7.79E-03 | 18.0 | 8.3E+04 | 1.7E-03 | 5.3 |
| | | **5.83E+05** | 3.55E+04 | **5.81E-03** | 3.17E-06 | 10.0 | | | | | | |
| | | **2.29E+05** | 1.69E+04 | **6.43E-03** | 1.05E-05 | 28.1 | | | | | | |
| | | **2.96E+05** | 1.90E+04 | **7.80E-03** | 2.80E-04 | 26.4 | | | | | | |
| | | **6.65E+05** | 1.63E+04 | **1.43E-02** | 1.78E-04 | 21.5 | | | | | | |
| | CAP256 | **1.33E+05** | 6.46E+05 | **2.26E-04** | <1.00E-06 | 1.69 | 1.21E+05 | 6.61E-04 | 5.47 | 1.5E+04 | 4.7E-04 | 3.92 |
| | | **1.39E+05** | 1.13E+06 | **1.59E-03** | 1.59E-03 | 11.5 | | | | | | |
| | | **9.07E+04** | 6.84E+05 | **1.65E-04** | <1.00E-06 | 1.82 | | | | | | |
| | JRFL | **9.19E+04** | 1.00E+06 | <1.00E-06 | **<1.00E-06** | 0.011 | 9.84E+04 | <1.00E-06 | 0.010 | 4.4E+03 | N/A | N/A |
| | | **1.07E+05** | 1.36E+06 | **<1.00E-06** | <1.00E-06 | 0.009 | | | | | | |
| | | **9.64E+04** | 3.43E+04 | <1.00E-06 | **<1.00E-06** | 0.010 | | | | | | |
| | BG505 GT1.2 | **8.27E+04** | 6.81E+05 | <1.00E-06 | **<1.00E-06** | 0.012 | 7.59E+04 | <1.00E-06 | 0.013 | 3.4E+03 | N/A | N/A |
| | | **7.26E+04** | 6.16E+05 | <1.00E-06 | **<1.00E-06** | 0.014 | | | | | | |
| | | **7.23E+04** | 6.12E+05 | <1.00E-06 | **<1.00E-06** | 0.014 | | | | | | |
| 2G12 IgG BCR | CH848 v4.1 DT | **3.17E+06** | 8.02E+04 | **1.80E-02** | 4.09E-04 | 5.68 | 3.21E+06 | 1.70E-02 | 5.30 | 1.7E+05 | 2.5E-03 | 0.81 |
| | | **2.94E+06** | 9.03E+04 | **1.24E-02** | 6.80E-08 | 4.21 | | | | | | |
| | | **3.53E+06** | 7.98E+04 | **2.07E-02** | 3.04E-04 | 5.86 | | | | | | |
| | CH505TF v4.1 | **7.42E+06** | 7.96E+03 | **5.74E-02** | 4.64E-04 | 7.73 | 7.18E+06 | 5.32E-02 | 7.40 | 8.6E+05 | 9.4E-03 | 1.58 |
| | | **8.55E+06** | 9.04E+03 | **6.69E-02** | 4.04E-04 | 7.83 | | | | | | |
| | | **5.58E+06** | 8.79E+04 | **3.53E-02** | 3.65E-04 | 6.32 | | | | | | |
| | CAP256 | **6.83E+05** | 3.25E+04 | <1.00E-06 | **1.29E-03** | 1.88 | 8.83E+05 | 1.02E-03 | 1.15 | 2.1E+05 | 2.9E-04 | 0.43 |
| | | **6.70E+05** | 2.43E+04 | <1.00E-06 | **1.33E-03** | 1.99 | | | | | | |
| | | **1.30E+06** | 3.00E+05 | <1.00E-06 | **4.35E-04** | 0.335 | | | | | | |
| | JRFL | **6.85E+05** | 1.49E+05 | **3.31E-04** | <1.00E-06 | 0.483 | 8.42E+05 | 1.61E-04 | 0.192 | 1.7E+05 | 8.6E-05 | 0.109 |
| | | **6.55E+05** | 1.48E+05 | <1.00E-06 | **9.76E-05** | 0.149 | | | | | | |
| | | **1.19E+06** | 2.30E+05 | **5.58E-05** | <1.00E-06 | 0.047 | | | | | | |
| | BG505 GT1.2 | **6.77E+05** | 1.79E+05 | <1.00E-06 | **1.45E-06** | 0.002 | 8.09E+05 | 1.39E-05 | 0.017 | 7.0E+04 | 8.6E-06 | 0.011 |
| | | **8.34E+05** | 2.21E+05 | **3.04E-05** | <1.00E-06 | 0.036 | | | | | | |
| | | **9.15E+05** | 2.22E+05 | <1.00E-06 | **9.85E-06** | 0.011 | | | | | | |
| 2G12 IgM BCR | CH848 v4.1 DT | **3.12E+06** | 6.07E+04 | **2.81E-02** | 9.99E-04 | 9.02 | 3.67E+06 | 2.69E-02 | 7.33 | 4.2E+05 | 6.0E-03 | 1.83 |
| | | **3.39E+06** | 9.03E+04 | **1.60E-02** | 3.43E-06 | 4.72 | | | | | | |
| | | **4.50E+06** | 6.55E+04 | **3.66E-02** | 3.92E-04 | 8.13 | | | | | | |
| | CH505TF v4.1 | **7.50E+06** | 1.92E+04 | **4.81E-02** | 2.59E-04 | 6.41 | 8.34E+06 | 5.96E-02 | 7.15 | 6.1E+05 | 6.4E-03 | 0.93 |
| | | **9.52E+06** | 4.56E+04 | **6.05E-02** | 2.34E-04 | 6.36 | | | | | | |
| | | **8.01E+06** | 9.24E+04 | **7.03E-02** | 8.67E-04 | 8.78 | | | | | | |
| | CAP256 | **9.29E+05** | 3.21E+04 | <1.00E-06 | **1.62E-03** | 1.74 | 1.03E+06 | 1.60E-03 | 1.55 | 1.1E+05 | 1.2E-05 | 0.16 |
| | | **9.17E+05** | 3.81E+04 | <1.00E-06 | **1.60E-03** | 1.75 | | | | | | |
| | | **1.25E+06** | 2.58E+05 | <1.00E-06 | **1.58E-03** | 1.26 | | | | | | |
| | JRFL | **8.01E+05** | 1.67E+05 | <1.00E-06 | **7.15E-05** | 0.089 | 1.03E+06 | 5.96E-05 | 0.058 | 2.3E+05 | 1.1E-05 | 0.017 |
| | | **8.04E+05** | 1.67E+05 | <1.00E-06 | **6.93E-05** | 0.086 | | | | | | |
| | | **1.50E+06** | 2.89E+05 | **3.82E-05** | <1.00E-06 | 0.026 | | | | | | |
| | BG505 GT1.2 | **9.86E+05** | 2.91E+05 | **2.28E-05** | <1.00E-06 | 0.023 | 1.13E+06 | 5.70E-05 | 0.050 | 7.4E+04 | 2.2E-05 | 0.020 |
| | | **1.18E+06** | 3.45E+05 | **9.82E-05** | <1.00E-06 | 0.083 | | | | | | |
| | | **1.23E+06** | 3.17E+05 | 1.52E-06 | **5.00E-05** | 0.041 | | | | | | |

**Table S1. Kinetic Rates Measurements for 2G12 [Relates to Main Figure 2].** Table of kinetic rates values from curve fitting using the heterogenous ligand model for all 2G12 interactions. Both sets of on (ka1 and ka2) and off-rates (kd1 and kd2) are provided, and bolded values indicate selected rates for affinity ($K_D$) calculation. All on-rates ($k_a$) selected have their curve fitting component account for ≥ 35% of maximum RU measured at end of the association phase for final injection of titration. All off-rates ($k_d$) selected are the faster of the two. Off-rates that are near or below limit of quantification by the instrument are reported as <1.00E-06 to indicate dissociation rates that are irreversible. Mean values are plotted in Main Figure 2.

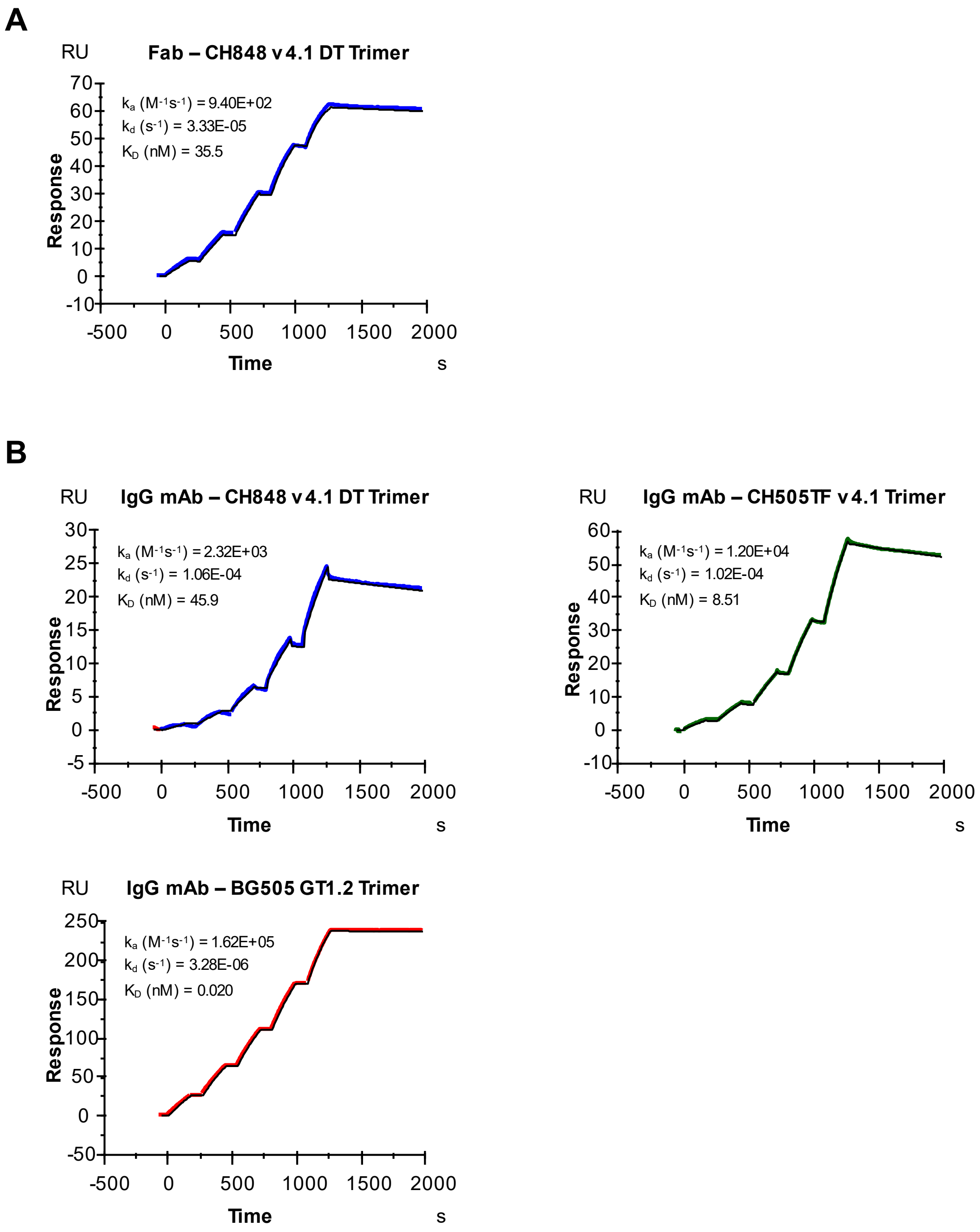

A
RU
Fab – CH848 v4.1 DT Trimer
k_a (M^-1 s^-1) = 9.40E+02
k_d (s^-1) = 3.33E-05
K_D (nM) = 35.5
Response
Time
s
B
RU
IgG mAb – CH848 v4.1 DT Trimer
k_a (M^-1 s^-1) = 2.32E+03
k_d (s^-1) = 1.06E-04
K_D (nM) = 45.9
Response
Time
s
RU
IgG mAb – CH505TF v4.1 Trimer
k_a (M^-1 s^-1) = 1.20E+04
k_d (s^-1) = 1.02E-04
K_D (nM) = 8.51
Response
Time
s
RU
IgG mAb – BG505 GT1.2 Trimer
k_a (M^-1 s^-1) = 1.62E+05
k_d (s^-1) = 3.28E-06
K_D (nM) = 0.020
Response
Time
s

C

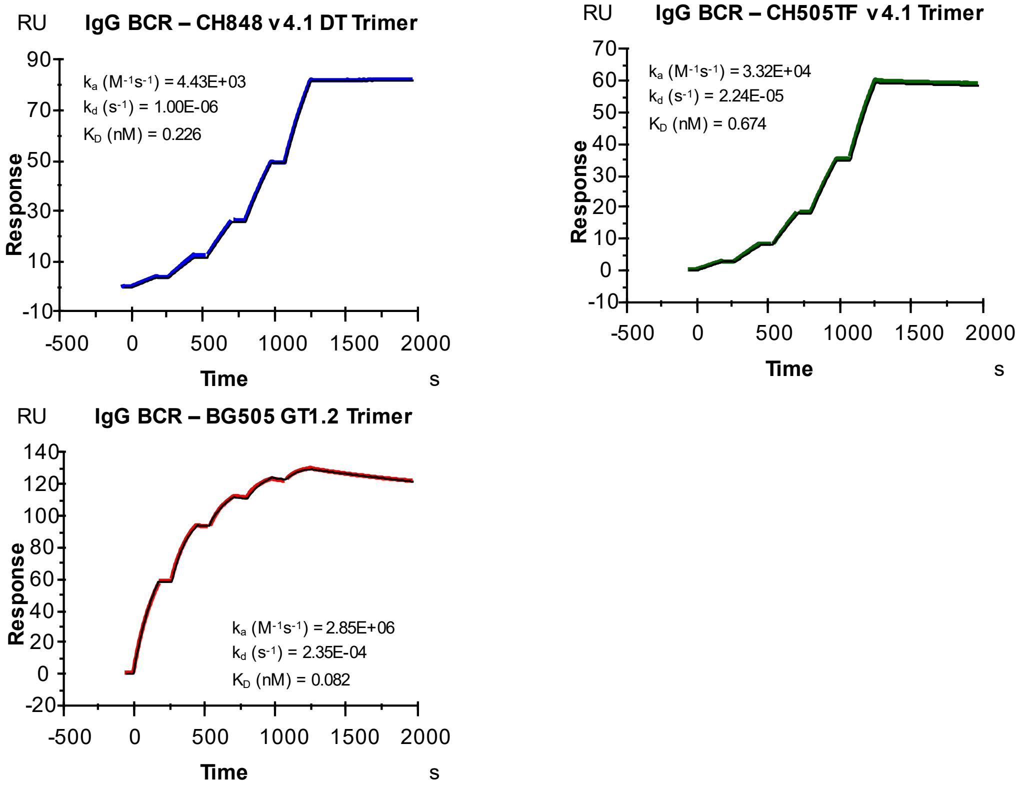

IgG BCR – CH848 v4.1 DT Trimer
RU
Response
Time
s
ka (M-1s-1) = 4.43E+03
kd (s-1) = 1.00E-06
KD (nM) = 0.226
IgG BCR – CH505TF v4.1 Trimer
ka (M-1s-1) = 3.32E+04
kd (s-1) = 2.24E-05
KD (nM) = 0.674
IgG BCR – BG505 GT1.2 Trimer
ka (M-1s-1) = 2.85E+06
kd (s-1) = 2.35E-04
KD (nM) = 0.082


D

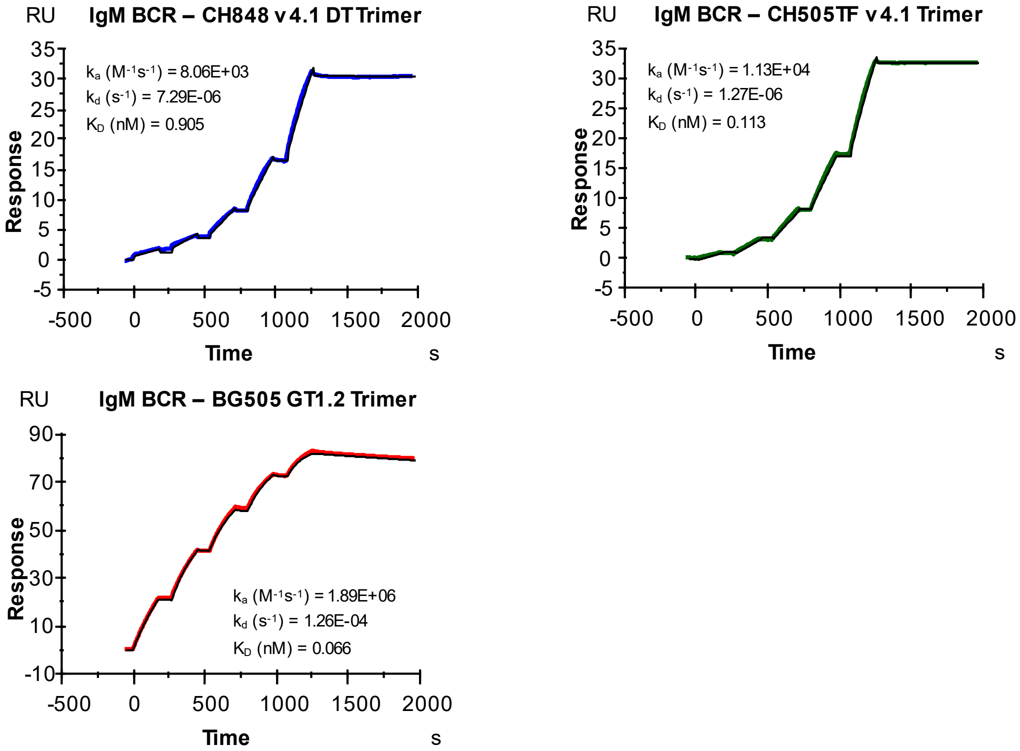

IgM BCR – CH848 v4.1 DT Trimer
RU
Response
Time
s
ka (M-1s-1) = 8.06E+03
kd (s-1) = 7.29E-06
KD (nM) = 0.905
IgM BCR – CH505TF v4.1 Trimer
ka (M-1s-1) = 1.13E+04
kd (s-1) = 1.27E-06
KD (nM) = 0.113
IgM BCR – BG505 GT1.2 Trimer
ka (M-1s-1) = 1.89E+06
kd (s-1) = 1.26E-04
KD (nM) = 0.066

**Fig. S4. SPR Curves and Fits for CH31. (A)** CH31 Fab flowed over biotinylated CH848 v4.1 DT Env trimer immobilized via streptavidin. **(B)** CH848 v4.1 DT, CH505TF v4.1, and BG505 GT1.2 Env trimer flowed over Anti-Human Fc captured CH31 IgG mAb. **(C)** Same Env trimers as in **B** flowed over captured purified CH31 IgG BCR. **(D)** Same Env trimers as in **B** flowed over captured purified CH31 IgM BCR. Surface Plasmon Resonance (SPR) was used to conduct titrations with Env trimers as described in methods section. CH31 Fab was diluted to concentrations ranging between 200 nM and 8000 nM. The following non-biotinylated Env trimers were used at the listed concentration ranges: CH505TF (2 – 200 nM), CH848 (10 – 1000 nM), GT1.2 (2 – 48 nM), CAP256 (1.5 – 48 nM), JRFL (2 – 48 nM). Heterogenous ligand model was used for biphasic curves. Plots shown are representative of at least three independent measurements.

| INTERACTION | | DATA FROM CURVE FITTING | | | | | MEAN | | | STANDARD ERROR | | |
|---|---|---|---|---|---|---|---|---|---|---|---|---|
| **Ligand:** | **Analyte:** | **ka1 (1/Ms)** | **ka2 (1/Ms)** | **kd1 (1/s)** | **kd2 (1/s)** | **KD (nM)** | **ka (1/Ms)** | **kd (1/s)** | **KD (nM)** | **ka (1/Ms)** | **kd (1/s)** | **KD (nM)** |
| Biotinylated CH848 v4.1 DT | CH31 Fab | **9.40E+02** | | **3.33E-05** | | 35.5 | | | | | | |
| | | **1.27E+03** | | **4.35E-05** | | 34.2 | 1.14E+03 | 3.89E-05 | 34.1 | 1.0E+02 | 3.0E-06 | 4.0 |
| | | **1.21E+03** | | **3.99E-05** | | 32.9 | | | | | | |
| CH31 IgG mAb | CH848 v4.1 DT | **2.32E+03** | | **1.06E-04** | | 45.9 | | | | | | |
| | | **1.62E+03** | | **1.10E-04** | | 67.6 | 1.92E+03 | 7.79E-05 | 40.7 | 2.1E+02 | 3.0E-05 | 16.3 |
| | | **1.81E+03** | | **1.76E-05** | | 9.75 | | | | | | |
| | CH505TF v4.1 | **1.20E+04** | | **1.02E-04** | | 8.51 | | | | | | |
| | | **1.20E+04** | | **9.21E-05** | | 7.68 | 1.17E+04 | 9.57E-05 | 8.19 | 2.8E+02 | 3.0E-06 | 0.32 |
| | | **1.11E+04** | | **9.33E-05** | | 8.38 | | | | | | |
| | BG505 GT1.2 | **1.62E+05** | 1.59E+06 | <1.00E-06 | **3.28E-06** | 0.0202 | | | | | | |
| | | **1.86E+05** | 1.87E+06 | <1.00E-06 | **5.40E-05** | 0.292 | 1.78E+05 | 3.67E-05 | 0.206 | 7.9E+03 | 1.7E-05 | 0.094 |
| | | **1.86E+05** | 1.87E+06 | <1.00E-06 | **5.28E-05** | 0.284 | | | | | | |
| CH31 IgG BCR | CH848 v4.1 DT | **4.43E+03** | | **<1.00E-06** | | 0.226 | | | | | | |
| | | **3.99E+03** | | **4.25E-06** | | 1.07 | 4.30E+03 | 2.08E-06 | 0.484 | 1.6E+02 | 1.1E-06 | 0.252 |
| | | **4.49E+03** | | **<1.00E-06** | | 0.223 | | | | | | |
| | CH505TF v4.1 | **3.32E+04** | | **2.24E-05** | | 0.674 | | | | | | |
| | | **2.55E+04** | | **2.99E-05** | | 1.17 | 2.80E+04 | 2.66E-05 | 0.949 | 2.6E+03 | 2.2E-06 | 0.118 |
| | | **2.54E+04** | | **2.75E-05** | | 1.08 | | | | | | |
| | BG505 GT1.2 | 4.20E+05 | **2.85E+06** | **2.35E-04** | <1.00E-06 | 0.082 | | | | | | |
| | | 4.31E+05 | **2.85E+06** | **2.73E-04** | <1.00E-06 | 0.096 | 2.84E+06 | 2.63E-04 | 0.093 | 1.1E+04 | 1.5E-05 | 0.005 |
| | | 4.34E+05 | **2.82E+06** | **2.82E-04** | <1.00E-06 | 0.100 | | | | | | |
| CH31 IgM BCR | CH848 v4.1 DT | **8.06E+03** | | **7.29E-06** | | 0.905 | | | | | | |
| | | **7.61E+03** | | **1.65E-06** | | 0.216 | 7.24E+03 | 3.31E-06 | 0.457 | 6.1E+02 | 2.0E-06 | 0.278 |
| | | **6.06E+03** | | **1.00E-06** | | 0.165 | | | | | | |
| | CH505TF v4.1 | **9.44E+03** | | **1.15E-06** | | 0.122 | | | | | | |
| | | **1.13E+04** | | **1.27E-06** | | 0.113 | 1.44E+04 | 3.21E-06 | 0.223 | 4.0E+03 | 2.0E-06 | 0.153 |
| | | **2.24E+04** | | **7.20E-06** | | 0.322 | | | | | | |
| | BG505 GT1.2 | **1.30E+06** | 2.04E+05 | **1.32E-04** | <1.00E-06 | 0.101 | | | | | | |
| | | **1.89E+06** | 2.77E+05 | **1.26E-04** | <1.00E-06 | 0.0665 | 1.69E+06 | 1.30E-04 | 0.077 | 2.0E+05 | 2.2E-06 | 0.009 |
| | | **1.89E+06** | 2.73E+05 | **1.33E-04** | <1.00E-06 | 0.0704 | | | | | | |

**Table S2. Kinetic Rates Measurements for CH31.** Table of kinetic rates values from curve fitting for all CH31 interactions. Bolded values indicate selected rates for affinity ($K_D$) calculation. For the heterogenous ligand model, both sets of on and off-rates are provided. All on-rates ($k_a$) selected have their curve fitting component account for ≥ 40% of maximum RU measured at end of the association phase for final injection of titration. All off-rates ($k_d$) selected are the faster of the two. Off-rates that are near or below limit of quantification by the instrument are reported as <1.00E-06 to indicate dissociation rates that are irreversible. Mean values are plotted in Main Figure 2.

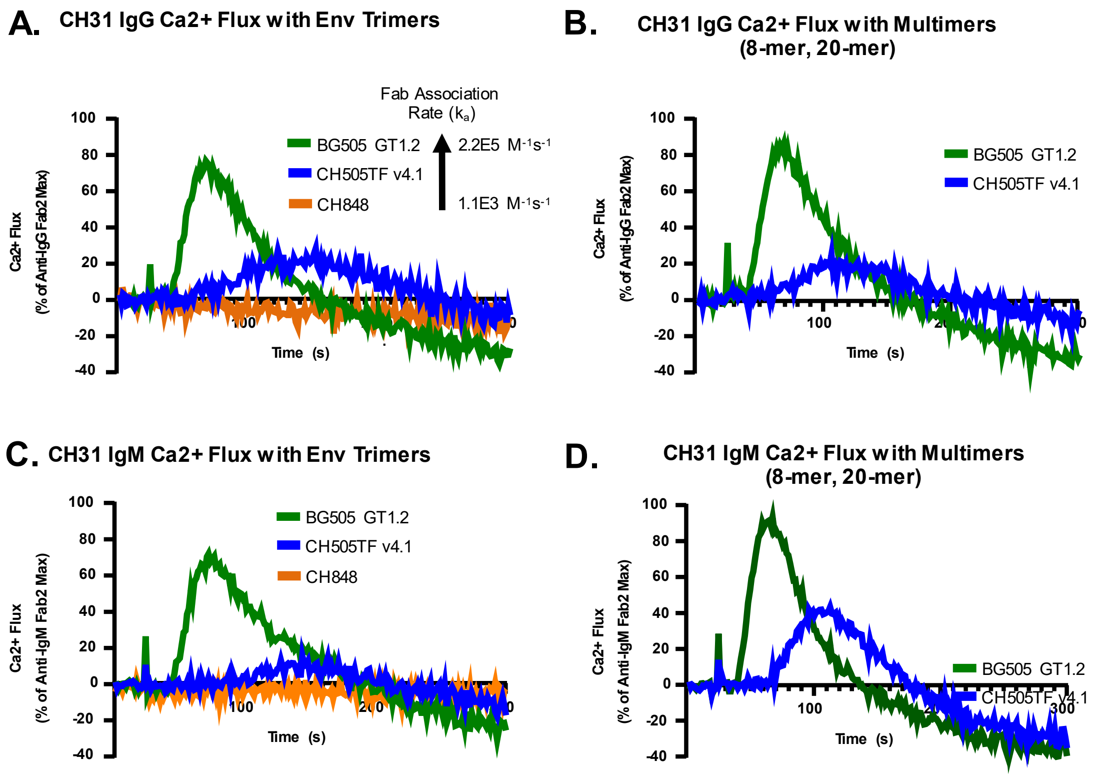
A. CH31 IgG Ca2+ Flux with Env Trimers
Fab Association Rate (ka)
2.2E5 M-1s-1
1.1E3 M-1s-1
BG505 GT1.2
CH505TF v4.1
CH848
Ca2+ Flux (% of Anti-IgG Fab2 Max)
Time (s)
B. CH31 IgG Ca2+ Flux with Multimers (8-mer, 20-mer)
BG505 GT1.2
CH505TF v4.1
Ca2+ Flux (% of Anti-IgG Fab2 Max)
Time (s)
C. CH31 IgM Ca2+ Flux with Env Trimers
BG505 GT1.2
CH505TF v4.1
CH848
Ca2+ Flux (% of Anti-IgM Fab2 Max)
Time (s)
D. CH31 IgM Ca2+ Flux with Multimers (8-mer, 20-mer)
BG505 GT1.2
CH505TF v4.1
Ca2+ Flux (% of Anti-IgM Fab2 Max)
Time (s)

**Fig. S5. Activation of Ramos Cells expressing IgM or IgG BCRs with specificity of CH31 to Env trimers and trimer multimers. (A)** Ca2+ flux responses induced in CH31 IgG Ramos cells by Env trimer proteins BG505 GT1.2 (green), CH505TFv4.1 (blue), and CH848 (orange) at 30nM. **(B)** Ca2+ responses induced in CH31 IgG Ramos cells by Env trimer multimers BG505 GT1.2 20-mer (green) and CH505TF v4.1 8-mer (blue) at 30nM. Ca2+ flux results for the CH31 IgG Ramos cells are presented as a percentage of the maximum anti-human IgG F(ab')$_2$ response and are an average of at least two measurements. Env trimers and their multimeric forms were prepared at the same per unit trimer concentrations for B cell activation (30nM). **(C)** Ca2+ flux responses induced in CH31 IgM Ramos cells by Env trimer proteins BG505 GT1.2 (green), CH505TFv4.1 (blue), and CH848 (orange) at 30nM. **(D)** Ca2+ responses induced in CH31 IgM Ramos cells by Env trimer multimers BG505 GT1.2 20-mer (green) and CH505TF v4.1 8-mer (blue) at 30nM. CH31 IgM Ramos cells were treated with Env trimers or multimers identically to the CH31 IgG Ramos cells. Ca2+ flux results for the CH31 IgM Ramos cells are presented as a percentage of the maximum anti-human IgM F(ab')$_2$ response and are an average of at least two measurements. Both CH31 IgG and IgG Ramos cell lines were developed with the AID enzyme knocked out (AIDKO).

**A**

| Env Trimer | 2G12 IgG Max Ca2+ Flux | 2G12 IgM Max Ca2+ Flux | Fold Change |
|---|---|---|---|
| CH505TF v4.1 | 26.4 | 65.5 | 2.5 |
| CAP256 | 48.3 | 86.4 | 1.8 |
| BG505 GT1.2 | 48.1 | 100.2 | 2.1 |
| JRFL | 32.6 | 115.3 | 3.5 |

**B**

| | 2G12 IgG Max Ca2+ Flux | | |
|---|---|---|---|
| **Env** | **Trimer** | **Multimer** | **Fold Change** |
| CH505TF v4.1 | 26.4 | 54.1 | 2.0 |
| CAP256 | 48.3 | 74.3 | 1.5 |
| BG505 GT1.2 | 48.1 | 79.8 | 1.7 |

**Table S3. Comparison of the maximum calcium flux (Ca2+ Flux) responses and activation of Ramos cells expressing 2G12 IgM or IgG BCRS with Env trimers and multimers [Relates to Main Figure 3] (A)** Summary of the maximum Ca2+ flux responses induced in 2G12 IgM Ramos and 2G12 IgG Ramos cells by Env trimer proteins. Ca2+ flux maximum responses for both cell lines are presented as a % of the maximum anti-human IgM or IgG F(ab')$_2$ response and are an average of at least two measurements. A comparison of the responses is presented as a fold-change to show the increase in Ca2+ flux induced in the 2G12 IgM Ramos cells versus the IgG cells. **(B)** Summary of the maximum Ca2+ flux responses induced in 2G12 IgG Ramos cells by Env trimer proteins and multimers. Ca2+ flux maximum responses for both cell lines are presented as a % of the maximum anti-human IgG F(ab')$_2$ response and are an average of at least two measurements. A comparison of the responses is presented as a fold-change to show the increase in Ca2+ flux induced by the multimeric forms of Env trimers proteins.

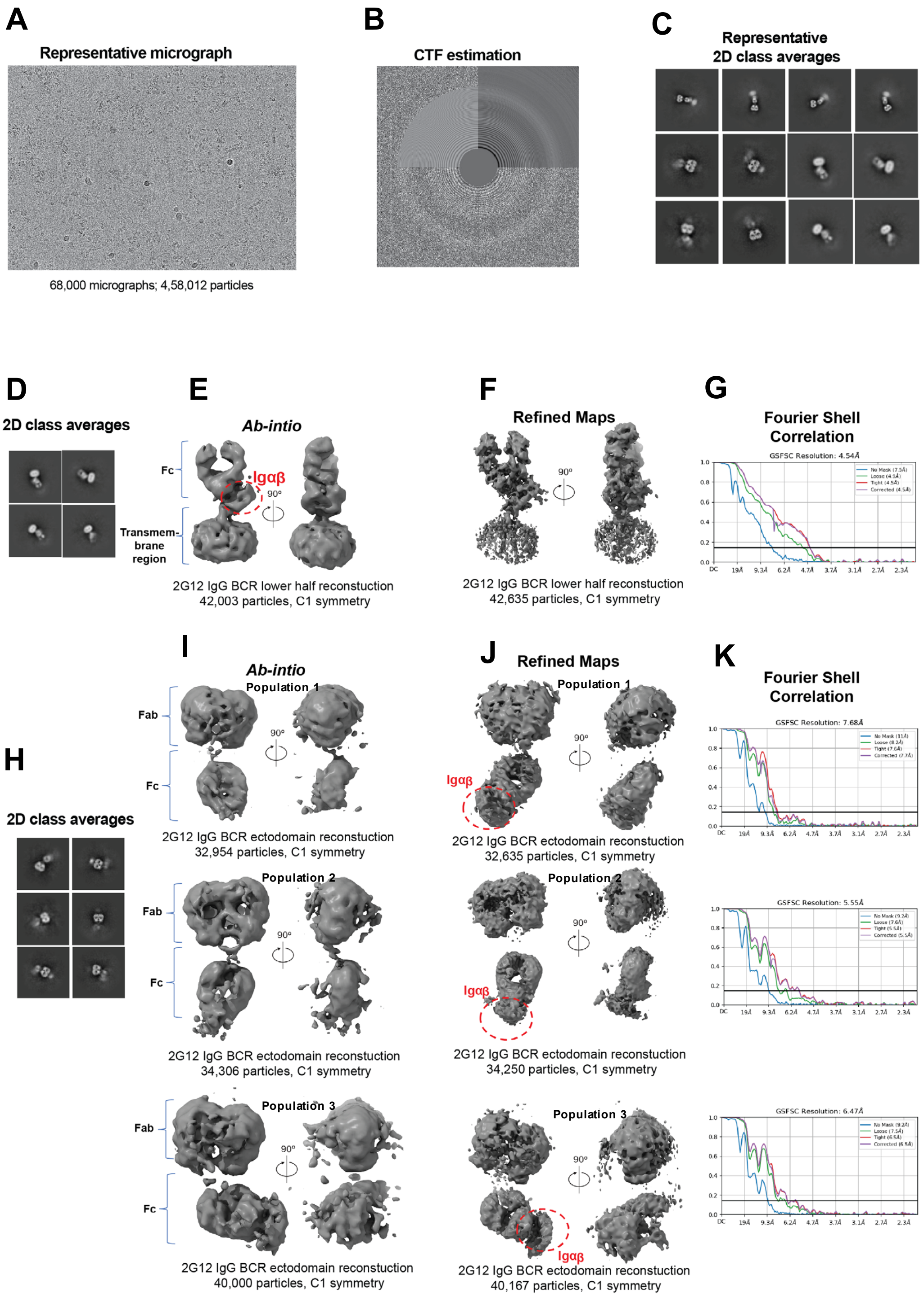
A
Representative micrograph
68,000 micrographs; 4,58,012 particles
B
CTF estimation
C
Representative
2D class averages
D
2D class averages
E
Ab-intio
Fc
Igαβ
90º
Transmem-
brane
region
2G12 IgG BCR lower half reconstuction
42,003 particles, C1 symmetry
F
Refined Maps
90º
2G12 IgG BCR lower half reconstuction
42,635 particles, C1 symmetry
G
Fourier Shell
Correlation
GSFSC Resolution: 4.54Å
H
2D class averages
I
Ab-intio
Population 1
Fab
Fc
90º
2G12 IgG BCR ectodomain reconstuction
32,954 particles, C1 symmetry
Population 2
Fab
Fc
90º
2G12 IgG BCR ectodomain reconstuction
34,306 particles, C1 symmetry
Population 3
Fab
Fc
90º
2G12 IgG BCR ectodomain reconstuction
40,000 particles, C1 symmetry
J
Refined Maps
Population 1
Igαβ
90º
2G12 IgG BCR ectodomain reconstuction
32,635 particles, C1 symmetry
Population 2
Igαβ
90º
2G12 IgG BCR ectodomain reconstuction
34,250 particles, C1 symmetry
Population 3
Igαβ
90º
2G12 IgG BCR ectodomain reconstuction
40,167 particles, C1 symmetry
K
Fourier Shell
Correlation
GSFSC Resolution: 7.68Å
GSFSC Resolution: 5.55Å
GSFSC Resolution: 6.47Å

**Fig. S6. Cryo-EM data processing for 2G12 IgG BCR**. **(A)** Representative micrograph. **(B)** Representative contrast transfer function (CTF) fit. **(C)** Representative 2D classes showing different views of BCR from cryo-EM dataset. Box size = 552.96 Å. **(D–G)** Clustering 2D class lacking Fab view together and its 3D reconstruction and refinement. **(D)** 2D class averages of particles lacking Fab region views. **(E)** Ab Initio reconstruction **(F)** Refined map. **(G)** Fourier shell correlation. A horizontal blue line in all FSC curves corresponds to a FSC value of 0.143. **(H–K)** Clustering 2D class lacking views for transmembrane region together and its 3D reconstruction and refinement. **(H)** 2D class averages. **(I)** Ab Intio reconstruction generating three different population classes representing differing conformations. **(J)** Refined maps for all three populations. **(K)** Fourier shell correlation. A horizontal blue line in all FSC curves corresponds to a FSC value of 0.143.

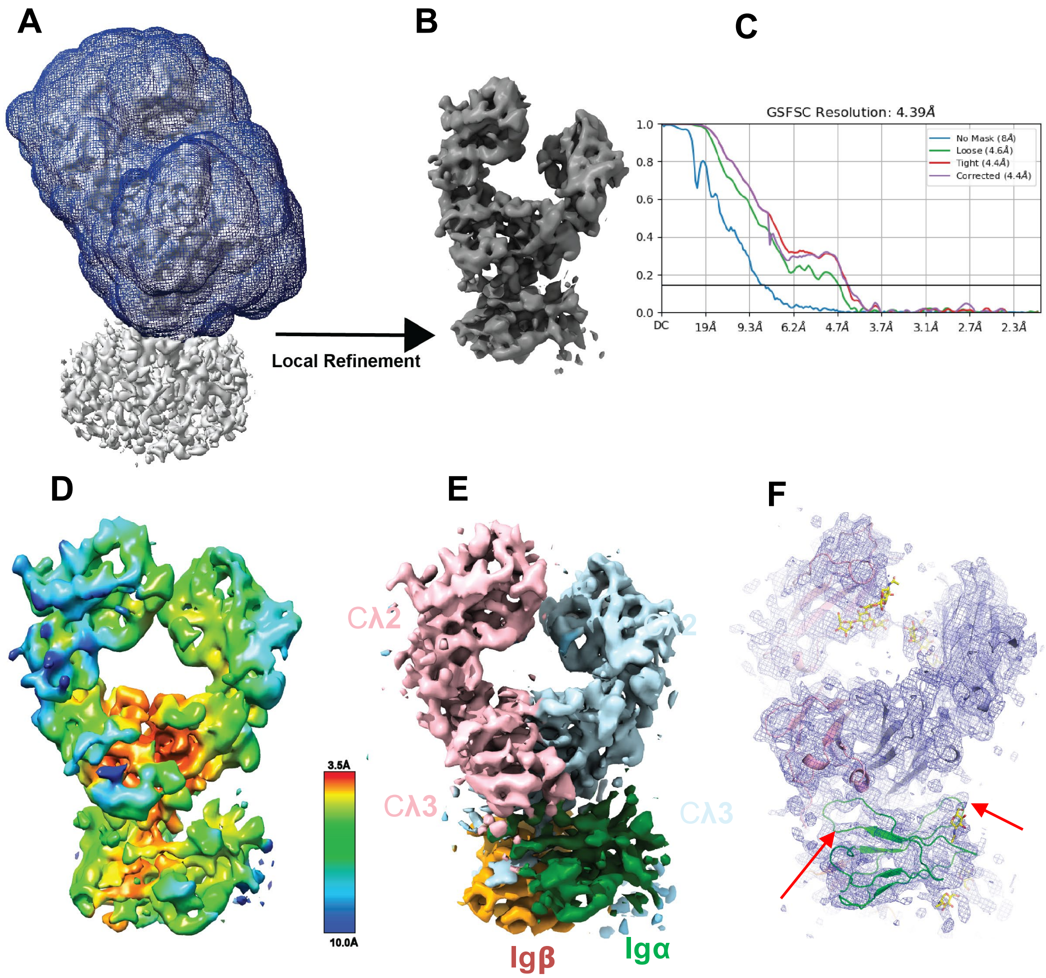


**Fig. S7.** Local resolution estimation and local refinement 2G12 IgG BCR. **(A)** Region selected for the local refinement by particle subtraction. **(B)** Locally refined map. **(C)** Fourier shell correlation. A horizontal blue line in all FSC curves corresponds to a FSC value of 0.143**. (D)** Refined region colored according to local resolution, where the red color represents best and blue showing worst resolution**. (E)** Refined map colored according to different BCRs structural components. **(F)** Fitted co-ordinate showing well defined density especially in the region forming the interacting shown by the red arrows.

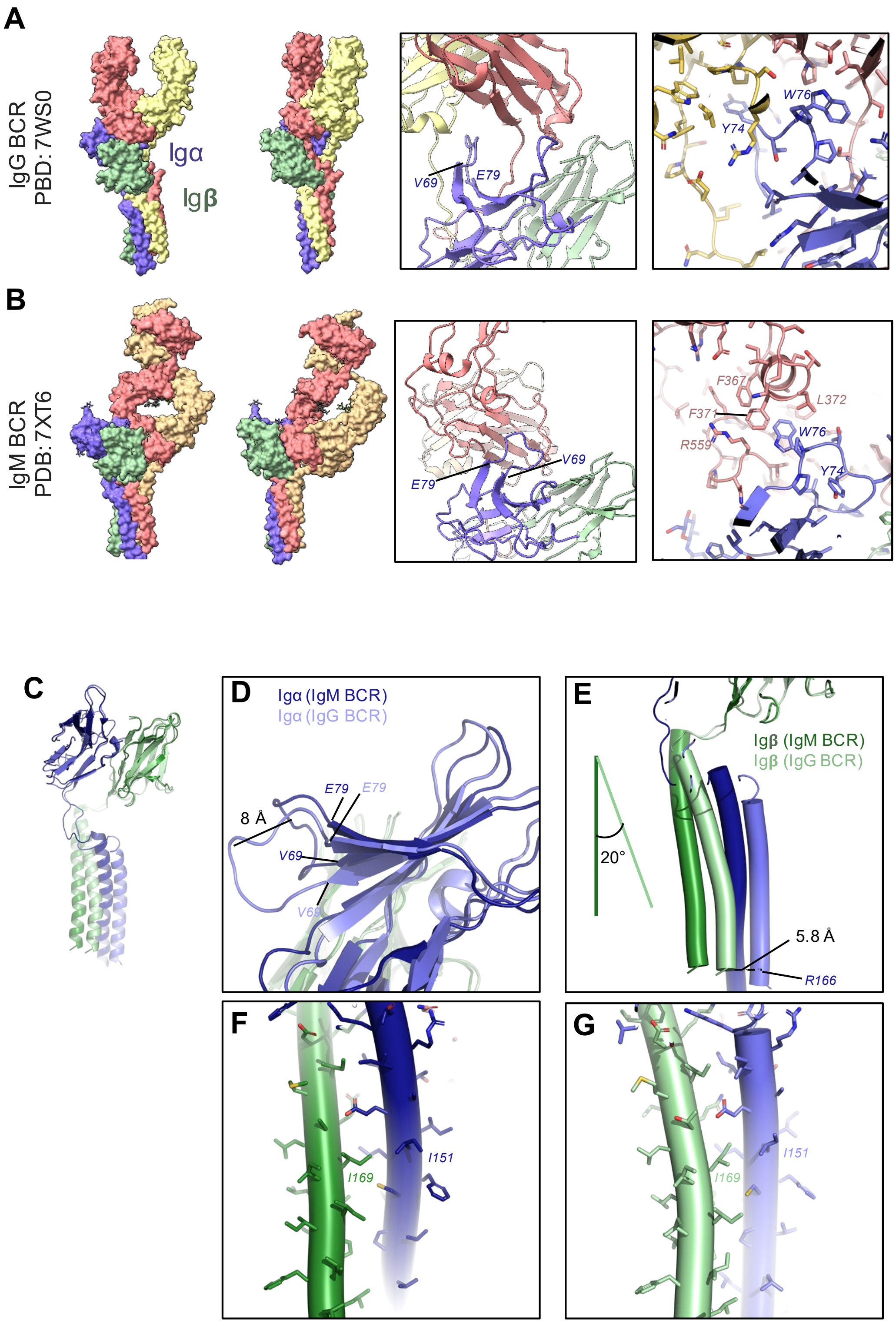

A
IgG BCR
PBD: 7WS0
Igα
Igβ
V69
E79
W76
Y74
B
IgM BCR
PDB: 7XT6
E79
V69
F367
L372
F371
W76
R559
Y74
C
D
Igα (IgM BCR)
Igα (IgG BCR)
E79
E79
8 Å
V69
V69
E
Igβ (IgM BCR)
Igβ (IgG BCR)
20°
5.8 Å
R166
F
I151
I169
G
I151
I169

**Fig. S8. Igα drives class-specific configuration of IgG versus IgM BCR.** **(A)** Left. Two views of IgG BCR shown in surface representation, with the Fc chains shown in salmon and yellow, Igα and Igβ colored blue and green, respectively. Middle. Zoomed-in view of interface between signaling components and Fc. Right. Zoomed-in view with side chains shown as sticks. **(B)** Same as in panel A. shown for IgM BCR. **(C)** Igα and Igβ chain of IgG and IgM BCRs superimposed with the Igβ ectodomain. **(D)** Zoomed-in view showing the loop spanning residues 69-79 of Igα that has an altered conformation between the IgG and IgM classes. **(E)** View of the TM helices, showing the differences between Igα and Igβ TM helix configurations. **(F)** TM helices for IgM BCR showing side chains as sticks. **(G)** TM helices for IgG BCR showing side chains as sticks.

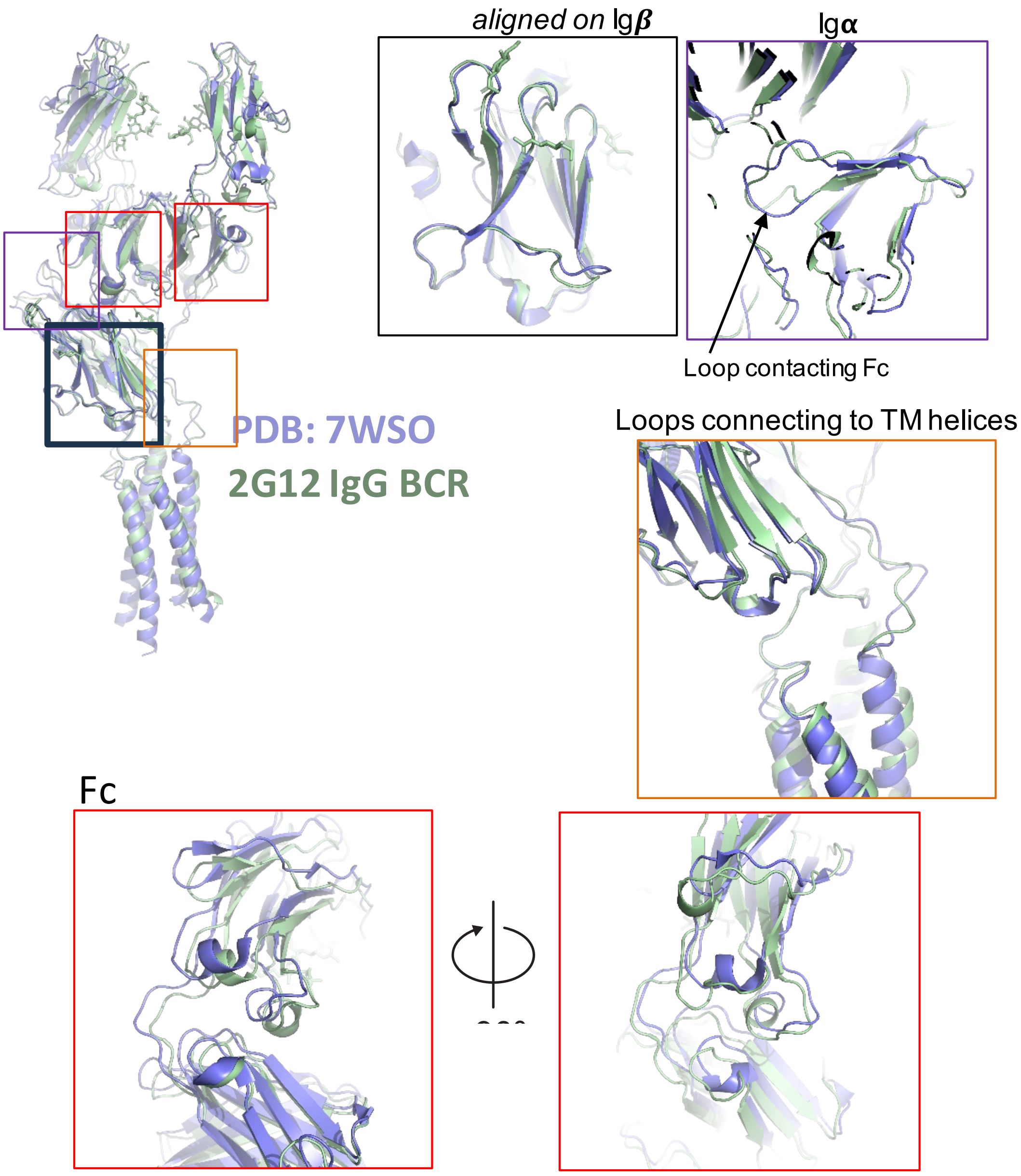


**Fig. S9. Comparison of 2G12 IgG BCR with 7WSO.** The two BCR structures (sans the Fab regions) were overlaid by their Igb domains. Zoomed in views of other domains are shown to visualize regions of flexibility in the BCR structure.

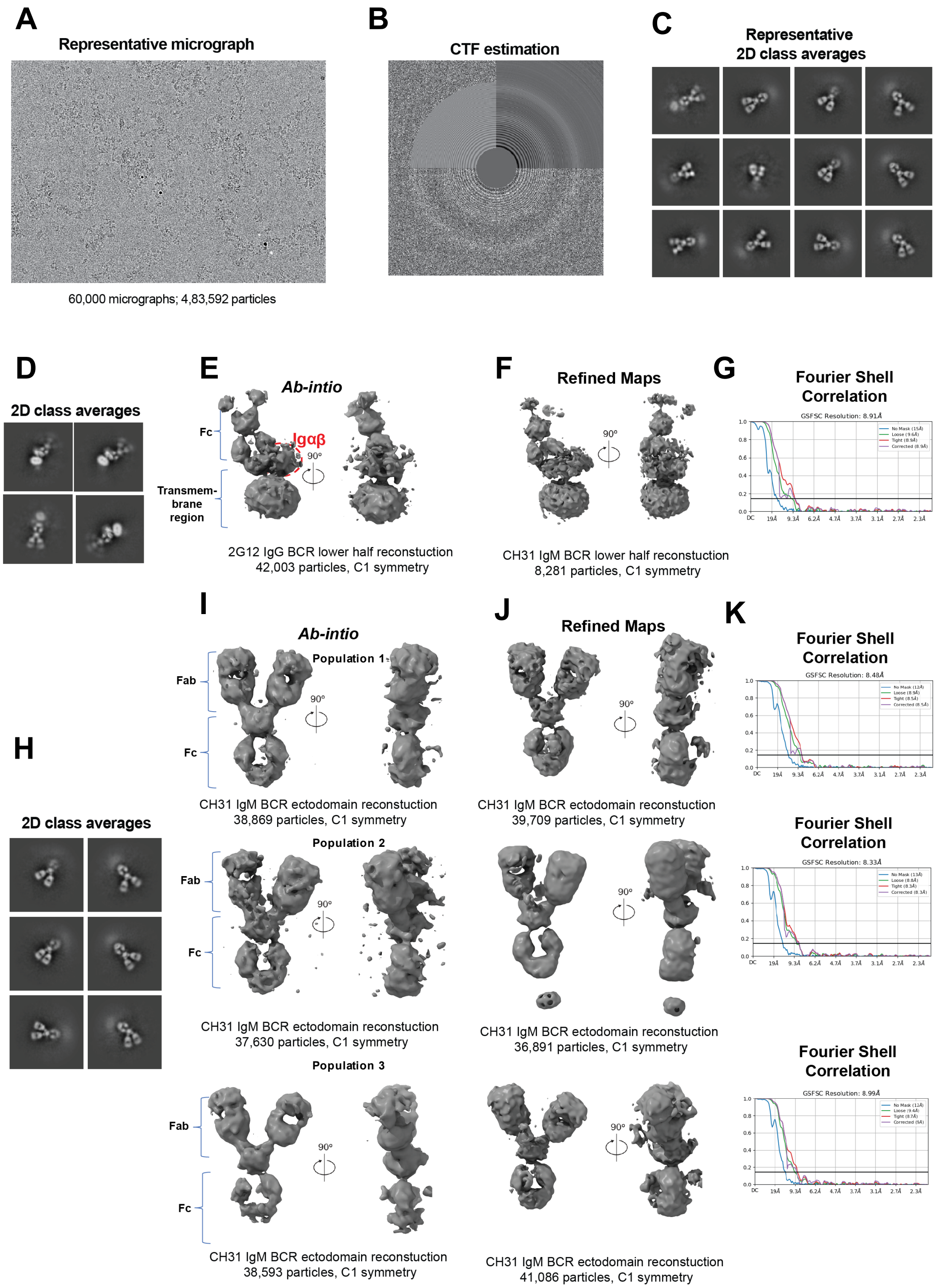
A
Representative micrograph
60,000 micrographs; 4,83,592 particles
B
CTF estimation
C
Representative
2D class averages
D
2D class averages
E
Ab-intio
Fc
Igαβ
90º
Transmem-
brane
region
2G12 IgG BCR lower half reconstuction
42,003 particles, C1 symmetry
F
Refined Maps
90º
CH31 IgM BCR lower half reconstuction
8,281 particles, C1 symmetry
G
Fourier Shell
Correlation
GSFSC Resolution: 8.91Å
H
2D class averages
I
Ab-intio
Population 1
Fab
Fc
90º
CH31 IgM BCR ectodomain reconstuction
38,869 particles, C1 symmetry
Population 2
Fab
Fc
90º
CH31 IgM BCR ectodomain reconstuction
37,630 particles, C1 symmetry
Population 3
Fab
Fc
90º
CH31 IgM BCR ectodomain reconstuction
38,593 particles, C1 symmetry
J
Refined Maps
90º
CH31 IgM BCR ectodomain reconstuction
39,709 particles, C1 symmetry
90º
CH31 IgM BCR ectodomain reconstuction
36,891 particles, C1 symmetry
90º
CH31 IgM BCR ectodomain reconstuction
41,086 particles, C1 symmetry
K
Fourier Shell
Correlation
GSFSC Resolution: 8.48Å
Fourier Shell
Correlation
GSFSC Resolution: 8.33Å
Fourier Shell
Correlation
GSFSC Resolution: 8.99Å

**Fig. S10. Cryo-EM data processing for CH31 IgM BCR**. **(A)** Representative micrograph**. (B)** Representative contrast transfer function (CTF) fit. **(C)** Representative 2D classes showing different views of BCR from cryo-EM dataset. Box size = 552.96 Å. **(D–G)** Clustering 2D class lacking Fab view together and its 3D reconstruction and refinement. **(D)** 2D class averages of particles lacking Fab region views. **(E)** Ab Initio reconstruction **(F)** Refined map**. (G)** Fourier shell correlation**.** A horizontal blue line in all FSC curves corresponds to a FSC value of 0.143. **(H–K)** Clustering 2D classes lacking views of transmembrane region together and its 3D reconstruction and refinement. **(H)** 2D class averages. **(I)** Ab Intio reconstruction generating three different population classes representing differing conformations. **(J)** Refined maps for all three populations**. (K)** Fourier shell correlation**.** A horizontal blue line in all FSC curves corresponds to a FSC value of 0.143.

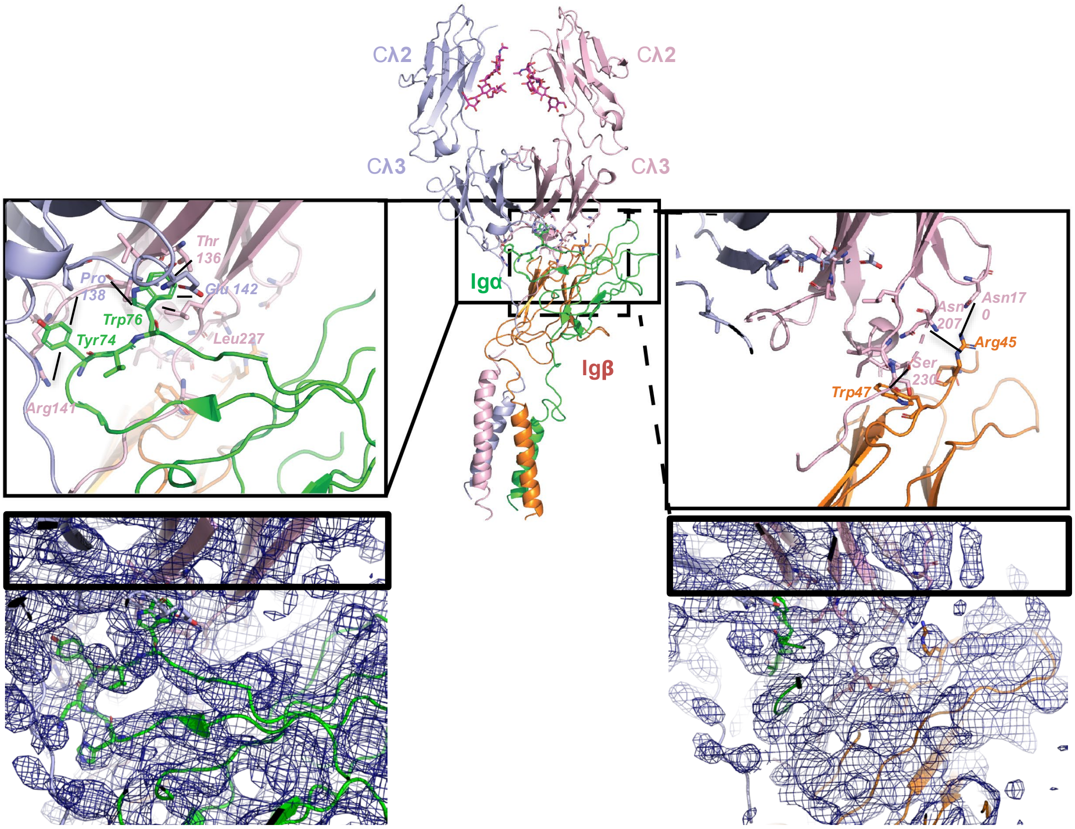


**Fig. S11. Interaction of co-receptors with Fc domains.** Left-Top. Zoomed-in view of interface between signaling components Igα and Cλ3 region of both of the heavy chains. Left-Bottom- Electron density map for the interacting residues. Right-Top. Zoomed-in view of interface between signaling components Ig**β** and Cλ3 region of one of the heavy chain. Right-Bottom. Electron density map for the interacting residues

| | Cryo-EM map of 2G12 IgG BCR lacking Fab region view | Locally Refined Cryo-EM map of 2G12 IgG BCR receptor lacking Fab region view | Cryo-EM map of 2G12 IgG BCR ectodomain, Population 1 | Cryo-EM map of 2G12 IgG BCR ectodomain, Population 2 | Cryo-EM map of 2G12 IgG BCR ectodomain, Population 3 | CH31 IgM BCR lacking Fab | cryo-EM map of CH31 IgM BCR representing immunoglobulin ectodomain population 1 | cryo-EM map of CH31 IgM BCR representing immunoglobulin ectodomain, population 2 | cryo-EM map of CH31 IgM BCR representing immunoglobulin ectodomain, population 3 |
|---|---|---|---|---|---|---|---|---|---|
| **PDB ID** | | 9E6G | | | | | | | |
| **EMDB ID** | EMD-47537 | EMD-47564 | EMD-47544 | EMD-47545 | EMD-47546 | EMD-47533 | EMD-47534 | EMD-47535 | EMD-47536 |
| **Data Collection and processing** | | | | | | | | | |
| Microscope | FEI Titan Krios | | | | | FEI Titan Krios | | | |
| Detector | Gatan K3 | | | | | Gatan K3 | | | |
| Magnification | 81000 | | | | | 81000 | | | |
| Voltage (kV) | 300 | | | | | 300 | | | |
| Electron exposure (e-/Å^2) | 59.1 | | | | | 59.1 | | | |
| Defocus Range (µm) | 3 to 2.3 | | | | | 3 to 2.3 | | | |
| Pixel size (Å) | 1.08 | | | | | 1.08 | | | |
| Micrographs collected | 68,000 | | | | | 60,000 | | | |
| Reconstruction software | cryoSPARC | | | | | cryoSPARC | | | |
| Symmetry imposed | C1 | C1 | C1 | C1 | C1 | C1 | C1 | C1 | C1 |
| Initial particle images (no.) | | | 4,83,592 | | | 987367 | | | |
| Final particle images (no.) | 42,635 | 42,635 | 32,659 | 34,250 | 40,167 | 8,281 | 39,709 | 36,891 | 41,086 |
| Map resolution (Å) | 4.54 | 4.39 | 7.68 | 5.5 | 6.47 | 8.91 | 8.48 | 8.33 | 8.99 |
| FSC threshold | 0.143 | 0.143 | 0.143 | 0.143 | 0.143 | 0.143 | 0.143 | 0.143 | 0.143 |
| **Refinement** | | | | | | | | | |
| Model resolution (Å) | | 4.56 | | | | | | | |
| FSC threshold | | 0.5 | | | | | | | |
| **Model composition** | | | | | | | | | |
| Nonhydrogen atoms | | 5143 | | | | | | | |
| Protein residues | | 618 | | | | | | | |
| **R.M.S. deviations** | | | | | | | | | |
| Bond lengths (Å) | | 0.009 | | | | | | | |
| Bond angles (°) | | 1.643 | | | | | | | |
| **Validation** | | | | | | | | | |
| MolProbity score | | 1.86 | | | | | | | |
| Clashscore | | 6.14 | | | | | | | |
| Favored rotamers (%) | | 99.34 | | | | | | | |
| **Ramachandran plot** | | | | | | | | | |
| Favored regions (%) | | 90.79 | | | | | | | |
| Disallowed regions (%) | | 0 | | | | | | | |

**Table S4.** **2G12 IgG and CH31 IgM BCR Cryo-EM data collection and refinement statistics.**

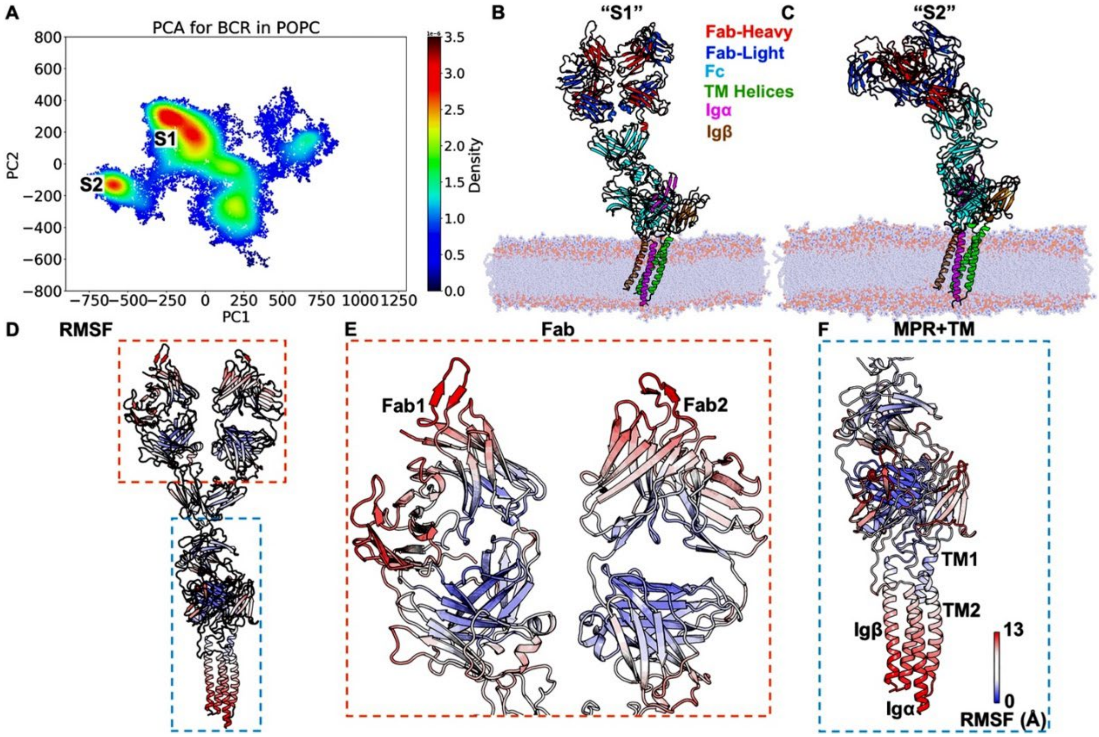

A
PCA for BCR in POPC
800
600
400
200
0
−200
−400
−600
−800
PC2
−750−500−250 0 250 500 750 1000 1250
PC1
S1
S2
3.5
3.0
2.5
2.0
1.5
1.0
0.5
0.0
Density
B
"S1"
Fab-Heavy
Fab-Light
Fc
TM Helices
Igα
Igβ
C
"S2"
D
RMSF
E
Fab
Fab1
Fab2
F
MPR+TM
TM1
TM2
Igβ
Igα
13
0
RMSF (Å)

**Fig. S12. Molecular Dynamics of free BCR in POPC lipid bilayer obtained from the MD simulations. (A)** Two-dimensional (2D) landscape of the distribution of the BCR conformations along the two primary principal components (PC) in the POPC lipid bilayer. A jet color scheme was used to show the distribution density of the BCR conformations. Two low-energy conformational states "S1"-"S2" were observed from the landscape based on their high densities. **(B)** The "S1" state, in which PC1 and PC2 coordinates were -225.7 and 305.8, respectively. **(C)** The "S2" state, in which PC1 and PC2 coordinates were -588.1 and -133.2, respectively. The Fab heavy chains were colored red, Fab light chains were colored blue, Fc domains were colored cyan, transmembrane helices of BCR were colored green, Ig$\alpha$ was colored magenta, and Ig$\beta$ was colored brown. The POPC molecules were colored light blue. **(D-F)** The flexibility of the BCR complex in POPC as measured by the root-mean-square fluctuations (RMSF). A color scale of blue (0.0) – white – red (13.0) is used to show the magnitudes of RMSFs.

**Movie. S1. Cryo-EM structures define dynamic positioning of 2G12 Fab in the BCR.** Movie showing different conformations of the Fab region derived from the flexibility of the linker (hinge) between the Fab and Fc region. The hinge region between the Fab and the Fc is not visible due to its flexible nature.

**Movie. S2. Dynamic of the CH31 BCR observed for atomistic molecular dynamics simulations**
